# New Swiss-knife activities of GroEL/Hsp60 proteins

**DOI:** 10.1101/2025.07.07.663466

**Authors:** Zhiyu Zhou, Dong Yang, Isaline Lambert, Corentin Decroo, Cyril Mascolo, Sophie-Luise Heidig, Tania Karasiewicz, Jean-François Flot, Martine Prévost, Ruddy Wattiez, Guy Vandenbussche, Véronique Fontaine

## Abstract

GroEL/Hsp60 chaperonins are key proteins that control cell metabolism, stress adaptation and survival. They usually form a tetradecameric structure that assists, coupled to ATP hydrolysis, 10% of all cellular protein folding. Using recombinant *E. coli*, human mitochondrial and *M. tuberculosis* chaperonins, we found that these proteins have thioesterase, esterase and even, for some of them, auto-acyltransferase activities. The smaller oligomers of Hsp60 and *M. tuberculosis* GroEL1 were more prone to use the long acyl carbon chain substrate palmitoyl-CoA compared to tetradecameric *E. coli* GroEL and Hsp60. Enzymatic competition and replacement of *M. tuberculosis* GroEL1 residues allow identifying Asp86 and Thr89 in the ATP-binding pocket and an additional Ser393 influencing the thioesterase activity. Additionally, *M. tuberculosis* GroEL1 might enhance palmitoylation of the PpsE protein, which plays a role in the phthiocerol dimycocerosate (PDIM) biosynthesis. This could explain at least partly the involvement of GroEL1 in PDIM biosynthesis and antibiotic resistance.

## INTRODUCTION

One of the most striking aspects in cell biology is the influence of environment on cellular mechanisms. Proteins that are involved in enzymatic activities must be timely synthesized and correctly folded so that their activity occur at the right place with the right partners. This is particularly important in stressful conditions. Chaperones, including heat shock proteins (HSPs), are known to facilitate protein folding. Chaperonins, some of the most important chaperones, are an evolutionary conserved group of proteins, composed of ∼ 55 kDa subunits assembled in 800-1000 kDa double rings^1^. Group I chaperonins, with *Escherichia coli* (*E. coli*) GroEL as an archetype, are found in all bacteria and in mitochondria (where it is called Hsp60) as well as in the chloroplasts of eukaryotes. They assist protein folding, thereby preventing protein aggregation^2^. These activities are dependent on ATP hydrolysis but also on their co-chaperone, GroES/Hsp10, which acts as a lid for each of their two heptameric rings stacked back-to-back^3^. Each of the 14 GroEL monomers consists of three domains: an apical domain involved in non-native protein and GroES binding through hydrophobic residues; an intermediate, highly flexible hinge domain able to encapsulate unfolded proteins; and an equatorial domain containing an ATP-binding site (DGTTT)^4^. Allosteric changes in the intermediate domain and large conformational changes inside the GroEL hydrophilic inner wall cage are driven by ATP binding in the equatorial domain. The 10 kDa GroES co-chaperone then binds to the GroEL apical domain, followed by ATP hydrolysis that triggers a change in polypeptide substrate folding^2,5^. By contrast, group II chaperonins found in archaea and in the cytosol of eukaryotes, (such as TCP-1 in human) also consist of two stacked rings, but they do not require a co-chaperone for their activity as they contain a built-in lid^6^. Group I and Group II chaperonins are distantly related, having probably diverged at the time of divergence of Bacteria from Archaea (including Eukaryotes) but it is still possible to align them together and draw them in a single all-encompassing tree (Figure S1).

Although in approximately 70% of bacteria (including *E. coli*) there is only one *gro*EL gene, two or three *gro*EL genes co-exist in the genomes of some lineages such as cyanobacteria^2^ and mycobacteria^7^. *Mycobacterium tuberculosis* contains two GroEL proteins, GroEL1 and GroEL2, also known as chaperonin 60, Cpn60.1 or Cpn60.2, or heat-shock protein 60, Hsp60-1 or Hsp60-2. GroEL1 shares 61% sequence identity with GroEL2, 52% sequence identity with its homologue *E. coli* GroEL and 38% sequence identity with human mitochondrial Hsp60 using the Clustal Omega alignment tools^8^. However, only *M. tuberculosis* GroEL2 is able to restore function in GroEL-defective *E. coli* mutants^9^. This could be partly explained by the fact that GroEL2 shows a stronger sequence identity with its homologue *E. coli* GroEL (59%). By contrast, *M. tuberculosis* GroEL1 plays a variety of essential roles under stress conditions that cannot be carried out by GroEL2^10–13^. These data suggest that GroEL proteins are “moonlighting proteins”, meaning that they assume different, unrelated functions in cells^14^. Unlike recombinant *E. coli* GroEL, recombinant *M. tuberculosis* GroEL1 and GroEL2 form smaller oligomers^9,15^, mainly monomers and dimers for GroEL1 and dimers and tetramers for GroEL2^16–19^. The ATPase activities of these two mycobacterial GroEL are very low (about 10 % for dimeric GroEL2 and about 6 % for GroEL1 compared to *E. coli* GroEL), not required for their folding and anti-aggregation activity and are not influenced by the presence of GroES^11,17,19^. A recent report mentioned that GroEL can hydrolyze a substrate of the β-galactosidase, the ortho-nitrophenyl β-galactoside^20^. Interestingly, acetylome analysis also revealed that *M. tuberculosis* GroEL1 and GroEL2, and *E. coli* GroEL, among others, are highly acetylated^21–23^. The physiological relevance of these post-translational modifications (PTMs) was however never studied, although PTMs have been shown to impact various cellular processes, both in prokaryotes and in eukaryotes. In bacteria, lysine acylation was reported to modify virulence^23,24^ and in eukaryotes, histone acetylation demonstrated a strong impact on gene expression.

GroEL/Hsp60 proteins have been targeted to develop therapeutic drugs against various diseases^25–29^. In the case of tuberculosis (TB), a severe infectious disease caused by a pathogenic mycobacterium from the *Mycobacterium tuberculosis* complex, GroEL1 has attracted interest. This disease, still causing 1.6 million deaths annually^30^, is difficult to treat, not least because of the *M. tuberculosis* impermeable cell wall. These bacteria have an external waxy mycomembrane, rich in mycolic acids, and an outer layer consisting in various non-covalently attached waxy lipids, including phenolglycolipids (PGL) and phthiocerol dimycocerosates (PDIM). PDIM lipids, long yβ-diols chain (phthiocerol) esterified with two long mycocerosic acids, are only present in pathogenic mycobacteria and involved in phagosomal escape, exit from host cells and drug resistance^12,31^. The mycobacterial GroEL1 protein is required for PDIM biosynthesis and for antibiotic resistance^12,32^. GroEL1 is also an essential virulence factor, required for *in vivo M. tuberculosis* persistence^32^. In infected lungs, *M. tuberculosis* can be detected intracellularly in macrophages, in granuloma lesions or extracellularly in necrotic lesions^33^ and counteracts environmental stresses thanks to GroEL1^10^.

Human mitochondrial Hsp60 acts as a classical protein-folding chaperonin involved in mitochondrion integrity but also exhibiting extra-mitochondrial functions that are not yet completely elucidated^25,34^. This protein was also targeted in drug-development as specific mutations reducing its ability to form oligomers have been associated with neurodegenerative diseases^25^ and its overexpression has been linked to inflammatory and autoimmune diseases as well as to various cancers^25,34–37^.

We report here the discovery that GroEL1 has a thioesterase activity. Its hydrolytic enzymatic activities were further investigated, uncovering that GroEL1 also had auto-acylation activities. The acetylated residues were identified, as well as those involved in these enzymatic activities. Furthermore, we verified whether *E. coli* GroEL and *human* Hsp60 also show hydrolytic activities. Finally, we studied whether GroEL1 could enhance PTM on other proteins, such as palmitoylation of a polyketide synthase protein, PpsE, involved in PDIM biosynthesis.

## Results

### GroEL2 is more evolutionary conserved than GroEL1 in terms of its structure and function, even though it is monocistronic

In *M. tuberculosis*, the structure of GroEL2 protein is quite similar to both HSP60 and TCP-1. It also shows the characteristic GMM C-terminal pattern in its sequence, found in archeal sequences as well as all across the bacterial part of the tree in Figure S1. This suggests that this GMM C-terminal pattern represents the ancestral state of GroEL proteins.

By contrast, GroEL1 is characterized by a histidine-rich C-terminus, which is rarely seen in the other clades represented on the phylogenetic tree in Figure S1. This highlights mycobacterial GroEL1’s evolutionary divergence from the ancestral GroEL pattern after following a gene duplication event that most probably occurred in the common ancestor of Actinobacteria, although GroEL1 retained its cistronic association with GroES following this duplication^7^.

Counterintuitively, GroEL2, although more divergent from archetypal GroEL in terms of operon structure, retained a function closer to archetypal GroEL, whereas GroEL1 retained the operon structure of the pre-duplication ancestor but acquired a divergent sequence and/or conformation responsible for additional activities, essential for stress survival^10–13,32,38^.

### GroEL/Hsp60 proteins have thioesterase and esterase activities

The thioesterase activity of TesA from *M. tuberculosis* was previously studied in our group^39^. Additional experiments showed that the cleavage of the palmitoyl-CoA thioester bond was increased when TesA was mixed with recombinant *M. tuberculosis* GroEL1 (Figure S2). This result prompted us to test for a possible thioesterase activity of GroEL1 itself, using ≥ 93 % (93-99%) purified GroEL1 (see the supplementary data and raw data on 10.5281/zenodo.13355298). As shown in Figure 1A, using palmitoyl-CoA as a substrate, GroEL1 exhibited thioesterase activity, with apparent kinetic parameters in the same range as for *M. tuberculosis* TesA activity^39^ (Figure 1A). In addition, GroEL1 showed an activity in a dose-dependent manner, the GroEL1 thioesterase reaction fitted to the apparent Michaelis-Menten model with a specific activity of 44.75 mU·mg^-1^ for palmitoyl-CoA. The kinetic parameters of the reaction included an apparent *V*_max_ (^app^*V*_max_) of (2.52 ± 0.25) × 10^-8^ M·s^-1^, an affinity for palmitoyl-CoA (^app^*K*_m_) of 25.95 ± 3.52 μM and a turnover number (^app^*k*_cat_) for palmitoyl-CoA of (2.52 ± 0.25) × 10^-3^ s^-1^.

**Figure 1.**
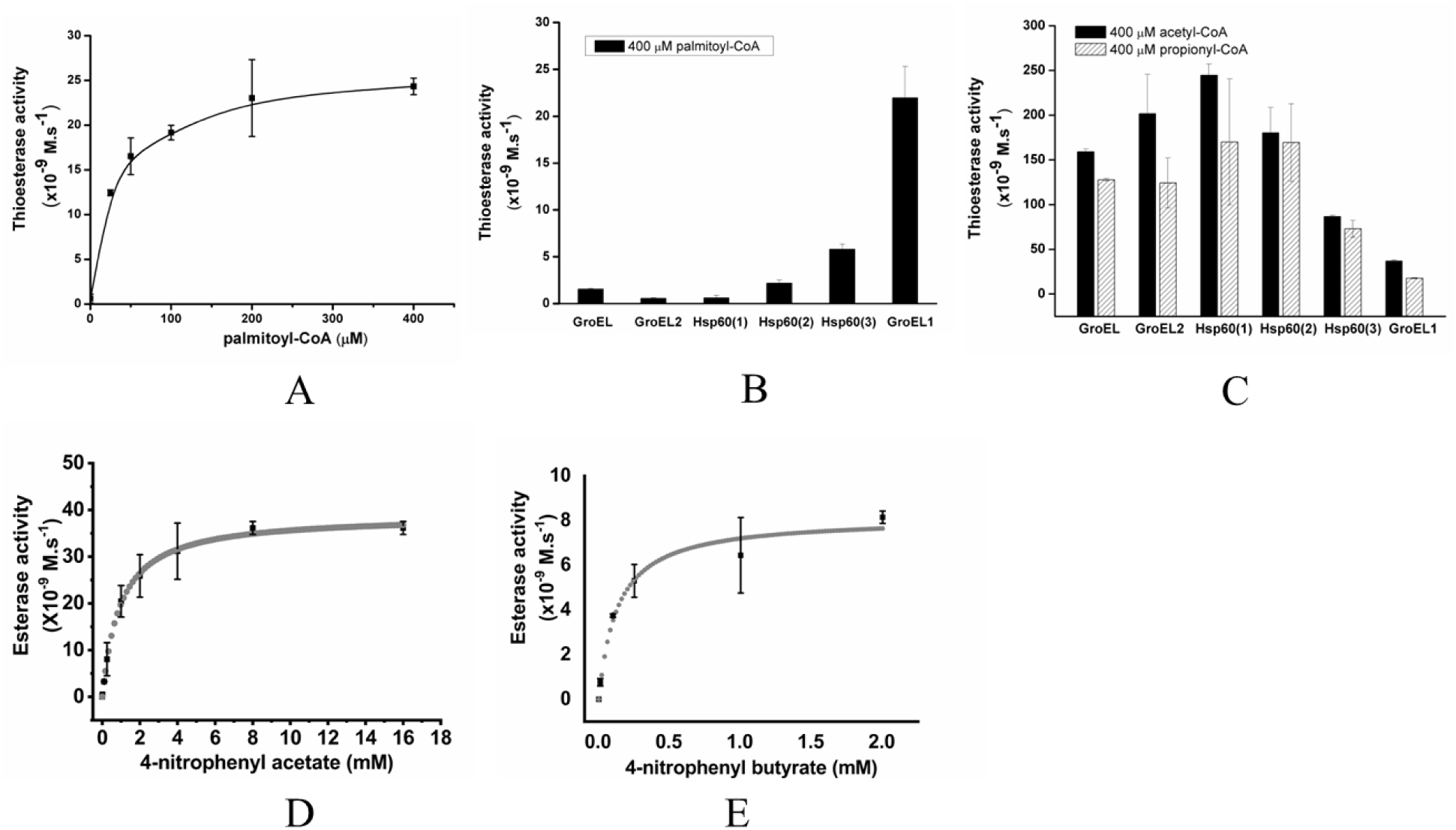
Thioesterase/esterase activities of recombinant proteins. (A) Dose-dependent thioesterase activity of recombinant GroEL1 in the presence of palmitoyl-CoA, (B) Comparison of different recombinant chaperonin thioesterase activity in the presence of 400 mM palmitoyl-CoA, (C) Recombinant chaperonin thioesterase activity in the presence of 400 mM acetyl-CoA and propionyl-CoA. GroEL1 esterase activity in the presence of 4-nitrophenyl acetate (D) or 4-nitrophenyl butyrate (E). The data were obtained from at least three independent experiments.

Several recombinant GroEL/Hsp60 chaperone proteins (Figures S3A, B and S4A, B) were additionally purified to test for thioesterase activity. Experimental molecular masses (without His-tag) of 61,345 Da for Hsp60, 56,169 Da for *M. tuberculosis* GroEL1, 57,620 Da for *E. coli* GroEL, 57,018 Da for *M. tuberculosis* GroEL2 were observed for these proteins using mass spectroscopy (MS), in agreement with their theoretical values (Figures S3C and S4C). Furthermore, different oligomeric forms of those recombinant proteins were detected by native-PAGE on 4-15 % gels (Figure S5). Recombinant Hsp60 presented various oligomeric forms that were further purified by size-exclusion chromatography (SEC) (Figures S4 and S5). When analyzing their thioesterase activities using palmitoyl-CoA as substrate, we observed that the recombinant Hsp60 (1), mainly present as two heptameric rings (tetradecamers), the recombinant Hsp60 (2) consisting mainly in heptamers, the tetradecameric recombinant *E. coli* GroEL and the various subunit forms of recombinant *M. tuberculosis* GroEL2 were inactive (Figure 1B). Only the lower subunit forms of recombinant Hsp60 (3) and the dimeric recombinant *M. tuberculosis* GroEL1 showed significant thioesterase activity on this substrate (Figure 1B). Nevertheless, we observed that all GroEL/Hsp60 chaperone proteins can cleave the thioester bond present in different short acyl carbon chain substrates, including acetyl-CoA and propionyl-CoA (Figure 1C).

Furthermore, as many thioesterase are also showing esterase activities^40^, the esterase activity of the recombinant *M. tuberculosis* GroEL1 was investigated using 4-nitrophenyl acetate (4-NPA) and 4-nitrophenyl butyrate (4-NPB) as substrates. GroEL1 displayed an esterase activity on 4-NPA and 4-NPB with an *^app^ V*max of (3.97 ± 0.39) × 10^-8^ M·s ^-1^ and (8.14 ± 0.28) × 10^-9^ M·s^-1^, respectively (Figure 1D and E). The apparent affinity for the 4-NPA and 4-NPB substrate (^app^*K*_m_) was 1.13 ± 0.74 mM and 0.18 ± 0.08 mM, respectively.

### Mycobacterial GroEL proteins are auto-acylated

Acetylation is a frequent modification that can occur either on the NH_3_^+^ group of the N-terminus of the protein (called Nt-acetylation) or on the side chain Nε group of lysine. In contrast to Nt-acetylation, lysine acetylation is a reversible PTM able to dynamically regulate protein functions both in eukaryotes and prokaryotes, being eventually critical for adaptation to stressful environmental conditions^41^. As such, acetylation is involved in virulence and pathogenicity. The human and *M. tuberculosis* acetylome studies have shown that Hsp60 and mycobacterial GroEL1 and GroEL2 proteins are among the most acetylated proteins^23,42^. Acetylation can be catalyzed by various acetyltransferase that can use acetyl-CoA as a precursor molecule. The discovery of the thioesterase activities of Hsp60 and GroEL1 prompted us to test the potential use of thioesterase reaction products for autoacyltransferase activities. Using propionyl-CoA as a substrate and a PTM protein enrichment strategy, based on immunoprecipitation with anti-propionyllysine antibodies, we identified the auto-propionylated residues of GroEL1 by MS/MS. Twelve lysine residues K3, K21, K243, K270, K275, K353, K362, K369, K388, K391, K423 and K524, according to the native GroEL1 sequence, were propionylated in the recombinant GroEL1 protein (Table 1A). The propionylation of the GroEL1 protein was verified on native *M. bovis* BCG GroEL1, which presents 100% sequence identity with the *M. tuberculosis* GroEL1. Of the 12 modified lysines detected in the recombinant protein, only 6 (K270, K275, K353, K369, K388 and K423) were found to be propionylated in the native form (Table 1B).

**Table 1.**
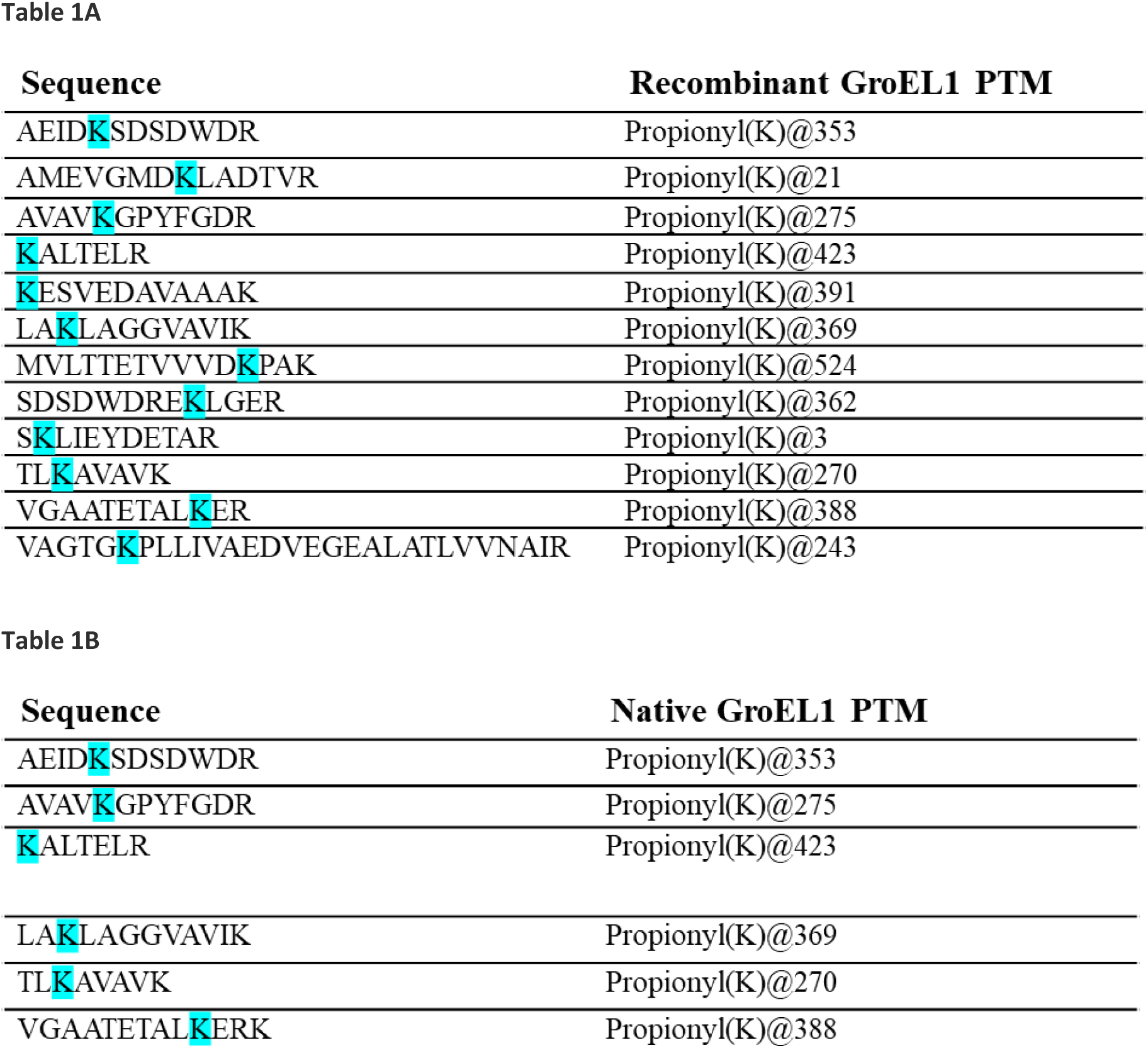
Propionylations on recombinant GroEL1 (A) and native *M. bovis* BCG GroEL1 (B).

Recombinant mycobacterial GroEL PTM was also investigated after reaction with 4-NPA and 4-NPB esterase substrates. After reaction with 4-NPA, a series of at least twelve peaks, with a Δ mass of 42 Da, were observed in the mass spectrum of GroEL1 (Figure S6A). Similarly, at least four peaks, with a Δmass of 70 Da were also observed in the presence of 4-NPB (Figure S6B). With GroEL2, up to seven peaks were observed in the presence of the 4-NPA, suggesting that recombinant GroEL2 was also able to be autoacetylated but in a slightly reduced manner compared to recombinant GroEL1 (Figure S6A). The acetylated residues of GroEL1 and GroEL2 after incubation with 4-NPA analyzed by MS/MS after trypsin/Lys-C or EndoGlu-C digestion allowed to locate the acetylation sites mostly on lysine residues of GroEL1 (K3, K21, K31, K79, K166, K345, K369, K378, K423, K468, K527) and GroEL2 (K3, K31, K369, K388, K391, K465). Another residue from GroEL1, S2, was also found to be acetylated (Figure S6C).

Finally, we investigated whether GroEL1 could be auto-palmitoylated. Due to difficulty to characterize such a hydrophobic PTM and in the absence of commercially available anti-palmitoyllysine antibodies, we tested auto-palmitoylation of recombinant mycobacterial GroEL using a fluorescent related palmitoyl-CoA. In these experimental conditions, recombinant GroEL1 was found to be palmitoylated which was not the case for GroEL2 (Figure 2A).

**Figure 2.**
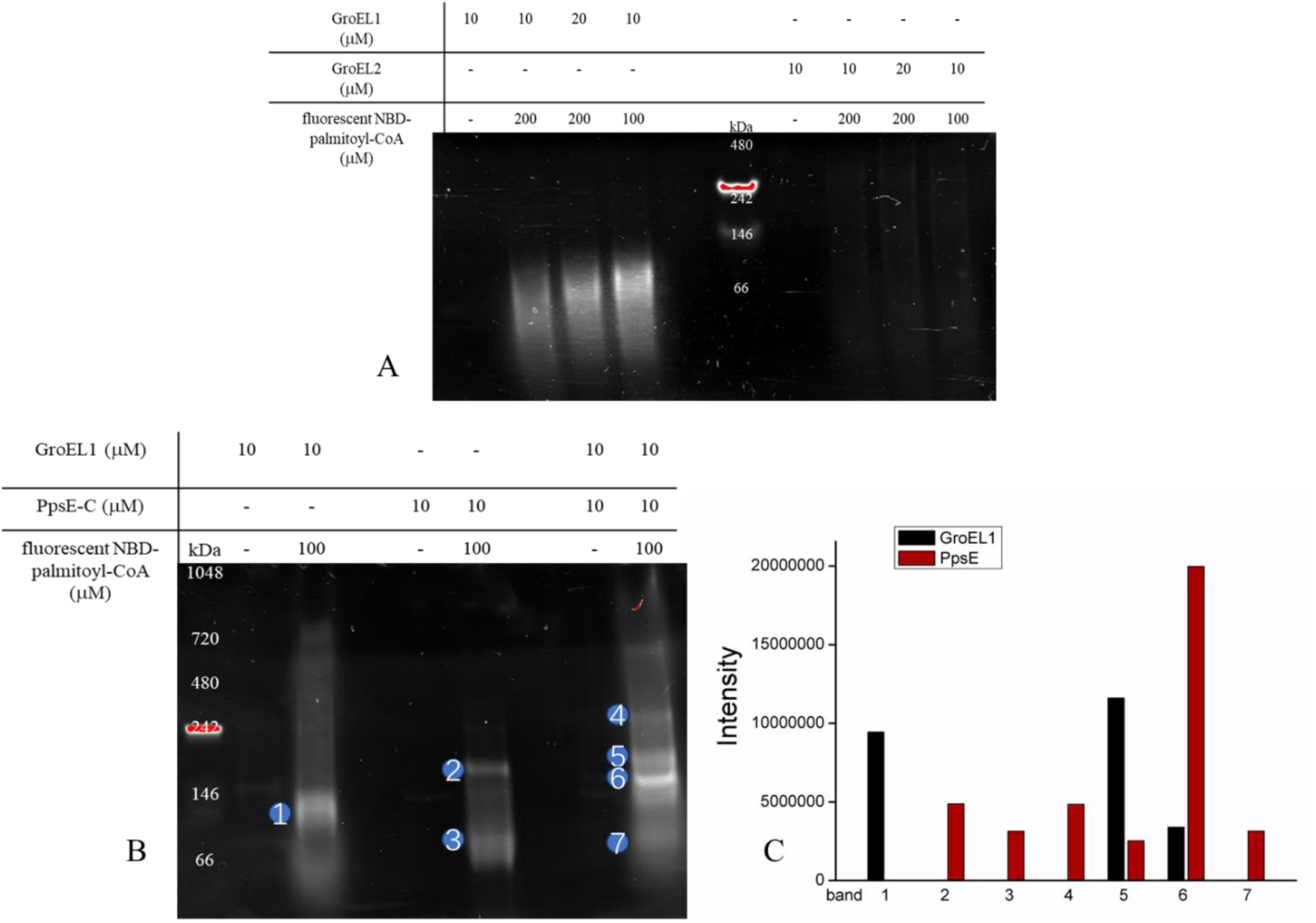
Detection by fluorescence of palmitoylated recombinant mycobacterial GroEL (A, B) and PpsE-C proteins (B) on 4-15 % native-polyacrylamide gel after 1h incubation at 37 °C in 100 mM sodium phosphate (pH 7.2) in the presence of eventually 10 mM recombinant mycobacterial GroEL and/or 10 mM PpsE-C protein and 100 mM fluorescent related palmitoyl-CoA. The data are representative of at least three independent experiments. LC-MS/MS identification analysis (C) of fluorescent digested proteins after elution from the native polyacrylamide gel (B).

### GroEL acyltransferase activity is modulating PpsE PTM

We previously shown that GroEL1 is required for PDIM biosynthesis^32^, for PDIM-related intrinsic resistance to vancomycin, for PDIM-related growth and for biofilm formation^10,12^, comparing wild-type (WT) and mutant Δ*cpn*60.1 (unable to produce GroEL1) *M. bovis* BCG strains^32^. Therefore, we investigated whether GroEL1 could have an impact on polyketide synthase (PKS) proteins, including the PpsA-E proteins involved in PDIM biosynthesis^43–45^, especially on potential PTM. First, we assessed the possibility that GroEL1 could bind PKS proteins by pull-down assay using WT and Δ*cpn*60.1 *M. bovis* BCG strain extracts. As GroEL1 can be retained on a Ni-column due to its natural Histidine-rich C-terminal region, we analyzed, by MS/MS, whether some proteins from the *M. bovis* BCG extracts could also be retained on this column, eventually through binding with GroEL1. Pks13, involved in mycolic acid biosynthesis and PpsA-E were jointly bound with GroEL1 to the Ni-column (Figure S7A). This suggests that GroEL1 could bind to these PpsA-E and Pks13 proteins. Nevertheless, it is worth mentioning that, in the absence of GroEL1, the production of *M. bovis* BCG PpsA-E proteins were reduced^10^. This probably explained the reduced amount of some high molecular weight (between 130 kDa and 250 kDa) proteins in the *M. bovis* BCG Δcpn60.1 protein extract compared to the WT *M. bovis* BCG strain protein extract (Figure S7B).

Secondly, we assessed the potential impact of GroEL1 on PTM of the carboxy half moiety of PpsE (PpsE-C), having approximately 60 kDa. When analyzing palmitoylation of recombinant GroEL1 and the soluble C-terminal half part of the PpsE protein (called here PpsE-C) in the presence of a fluorescent palmitoyl-CoA analogue, we not only observed GroEL1 auto-palmitoylation, as it could be expected from its thioesterase activity (Figure 2A), but also PpsE-C auto-palmitoylation. The PpsE-C auto-palmitoylation was approximately similar to the GroEL1 auto-palmitoylation. Surprisingly, when both proteins were incubated in the same reaction, palmitoylation fluorescence signal intensity was further increased (Figure 2B). LC/MSMS analysis of gel-eluted fluorescent proteins, after trypsin digestion, demonstrated an increased palmitoylation of PpsE-C in the presence of GroEL1 and even suggested that GroEL1 and PpsE could interact with each other (Figure 2C).

### GroEL1 thioesterase catalytic site and identification of functional residues

Having demonstrated the thioesterase activity of GroEL1, we sought to locate the site of this enzymatic activity and the residues likely to be functionally important. Palmitoyl-CoA consists in three chemical moieties: the hydrophobic palmitic acid chain, the (phospho)pantetheine and the adenosine tri-(or di-)phosphate (ATP/ADP) analogue (Figure 3A). In view of the analogy of the latter with ADP, we hypothesized that the ATP binding site, previously identified in the *E. coli* GroEL^46^, could interact with the ADP-like part of the acyl-CoA. As such, ATP would act as a competitive inhibitor of the GroEL1 thioesterase activity. As shown in Figure 3B, GroEL1 thioesterase activity is indeed inhibited by ATP, in a dose-dependent manner, especially in the presence of magnesium. As ATP and Mg are both required for enabling *E. coli* GroEL protein to form an oligomeric complex with GroES, an essential key step in the protein folding process^47^, the impact of ATP and Mg was therefore investigated on *M. tuberculosis* GroEL1 oligomerization. As observed on native polyacrylamide gel, in the presence of ATP and Mg (Figure S8A), the amount of dimeric GroEL1 was strongly reduced, although GroEL1 could still be detected in denaturing conditions. Moreover, the addition of 5 mM ATP and 10 mM MgSO_4_ also improved the stability of the GroEL1, as observed in thermal shift assay (TSA) (Figure S8B). ATP binding could therefore not only compete in the GroEL1 thioesterase activity, but it could also reduce thioesterase activity by affecting GroEL1 oligomerization.

**Figure 3.**
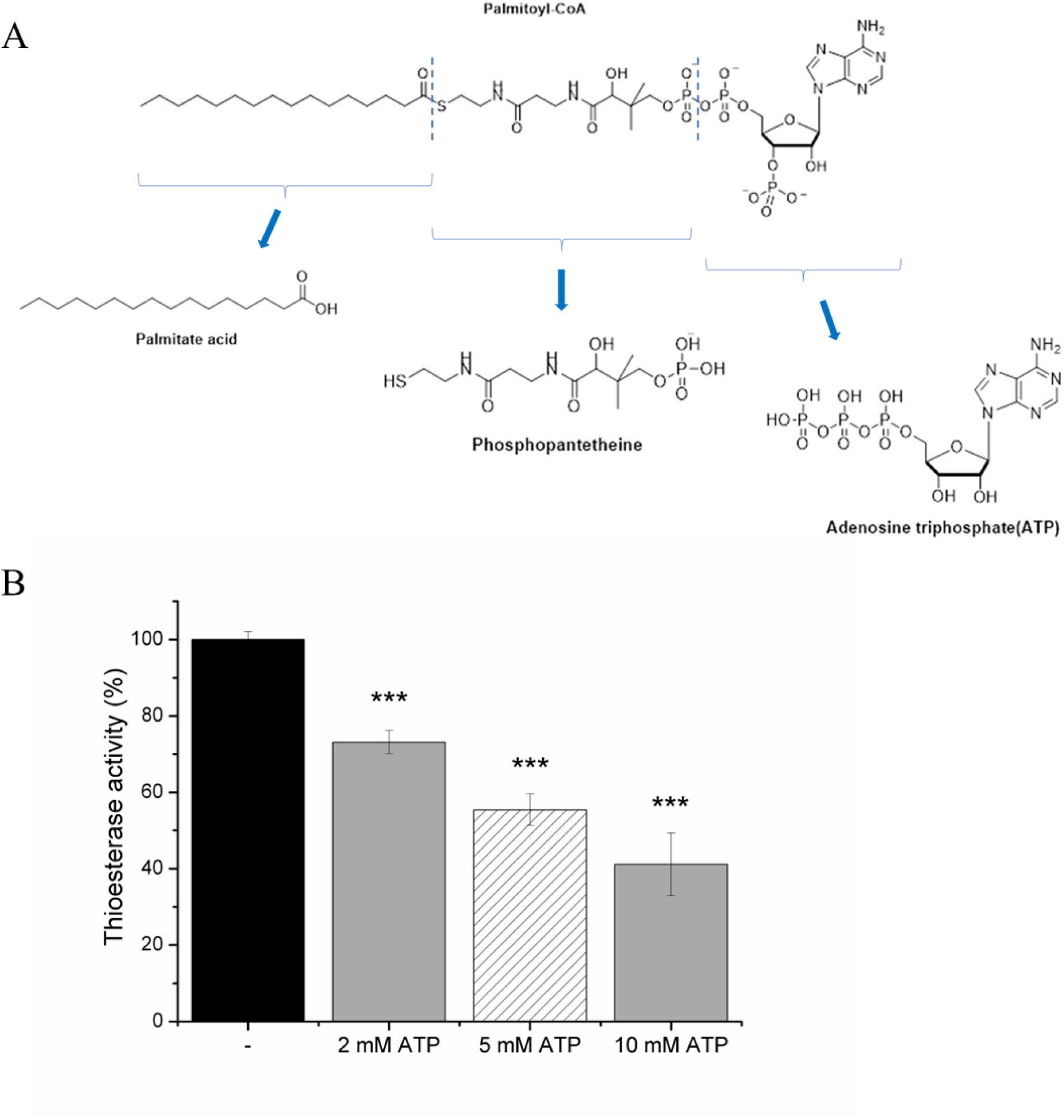
(A) Palmitoyl-CoA structure. (B) Dose-dependent impact of ATP on GroEL1 thioesterase activity. The activity was monitored by measuring the absorbance at 412 nm for 30 min at 37 °C. The data were obtained from at least three independent experiments. \*\*\**p* <0.001.

To further investigate the role of the ATP binding pocket in the GroEL1 thioesterase activity, the docking calculations were carried out. Firstly, to validate our procedure, the docking of AGS, an ATP analog, was performed in different modelled structures and compared to its crystalline position in the X-ray structure of the *E. coli* GroEL homologue. The AGS position closest to the experimental geometry was obtained for the docking in the GroEL1 3D model generated using as a template the *Xanthomonas oryzae* GroEL structure and features the phosphate groups and magnesium forming interactions with D86 or with the threonine triplet 88-90 (Figure 4A). In addition, the AI tool AlphaFill^48^, which is using information of experimentally determined binding pockets and ligands to identify and fill matching areas in predicted protein structures, also agrees for a potential ATP binding pocket 86DGTTT90 in the *M. tuberculosis* GroEL1 AlphaFold model (P9WPE9 from Uniprot).

**Figure 4.**
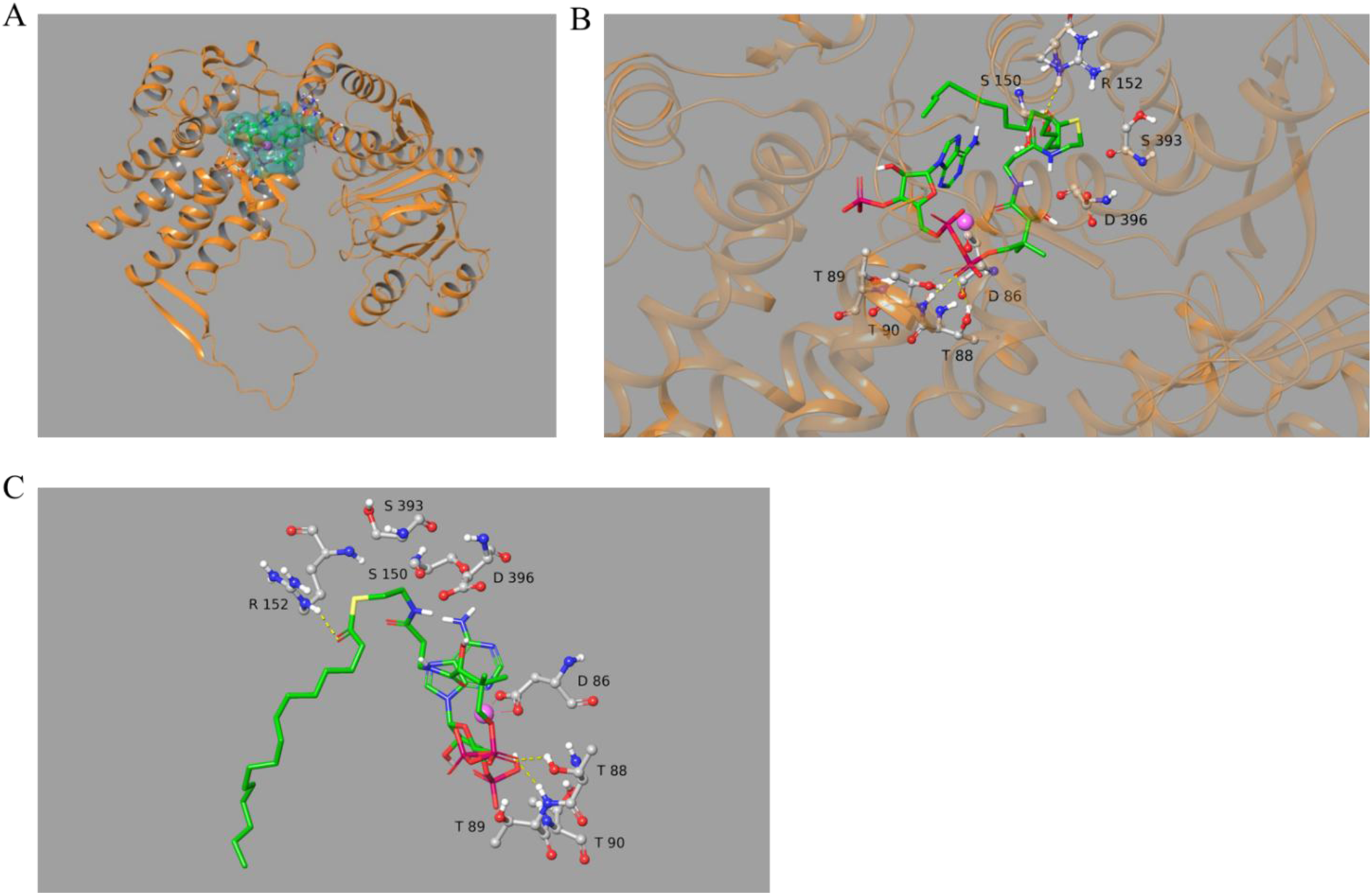
(A) Palmitoyl-CoA was docked in the ATP binding pocket of the model from *Xanthomonas oryzae* in the presence of magnesium. Details are shown in panel B and C.

Palmitoyl-CoA was then docked in the ATP binding pocket of all modeled structures. Docked positions were selected based on their similarity to the position of AGS (Figure 4A). In particular, the proximity of the phosphate groups to D86 or the 88-90 threonine triplet as with AGS was used as a criterion. On this basis, only the positions obtained from the docking in the structure modeled with *X. oryzae* GroEL in the presence of magnesium were selected (Figure 4B, 4C).

Analysis of the positions of the docked palmitoyl-CoA led to the selection of several residues based on their proximity to the thioester and phosphate groups, namely D86, T89, S150, D396, R152, S393, for their mutation in the cpn60.1 gene in order to change the corresponding residues into alanine. Kinetic of H/D exchanged analyzed by FTIR showed that the tertiary structure of the purified mutant proteins (Figures S9A and B) was close to that of the WT protein, with no significant change in global conformation (Figure S9C).

The mutations introduced in GroEL1 were mapped onto corresponding residues in other GroEL/Hsp60/TCP-1 proteins using the full MSA. MSA logos allow to assess residue conservation at the pinpointed positions, potentially highlighting functional implications of these residues (Table S4). Positions 116 and 119 in the MSA represent D86 and T89 in the highly conserved ATP-binding site motif DGTTT. Positions 187 and 201 in the MSA correspond in GroEL1 sequence to S150 and R152, respectively. The sequence logos reveal that, while uncommon, an alanine at these positions is occurring in natural GroEL/Hsp60 sequences. At position 201 the sequence conservation is indeed lower. MSA position 526 corresponds to S393 in GroEL1, an unusual residue as it is usually a highly conserved arginine residue at this position. Position 529 corresponds to D396, which is revealed to be another highly conserved residue.

The thioesterase activity of all mutant proteins was then tested with palmitoyl-CoA as substrate. Only the thioesterase activities of the D86A, T89A, R152A and S393A GroEL1 mutant proteins decreased significantly compared to that of the WT protein (WT) (Figure 5). The activity of the D86A GroEL1 was less than 20% compared to the WT GroEL1 activity. The T89A and the S393A mutant GroEL1 showed less than 40% and 50% activity, respectively, compared to the WT protein (Figure 5).

**Figure 5.**
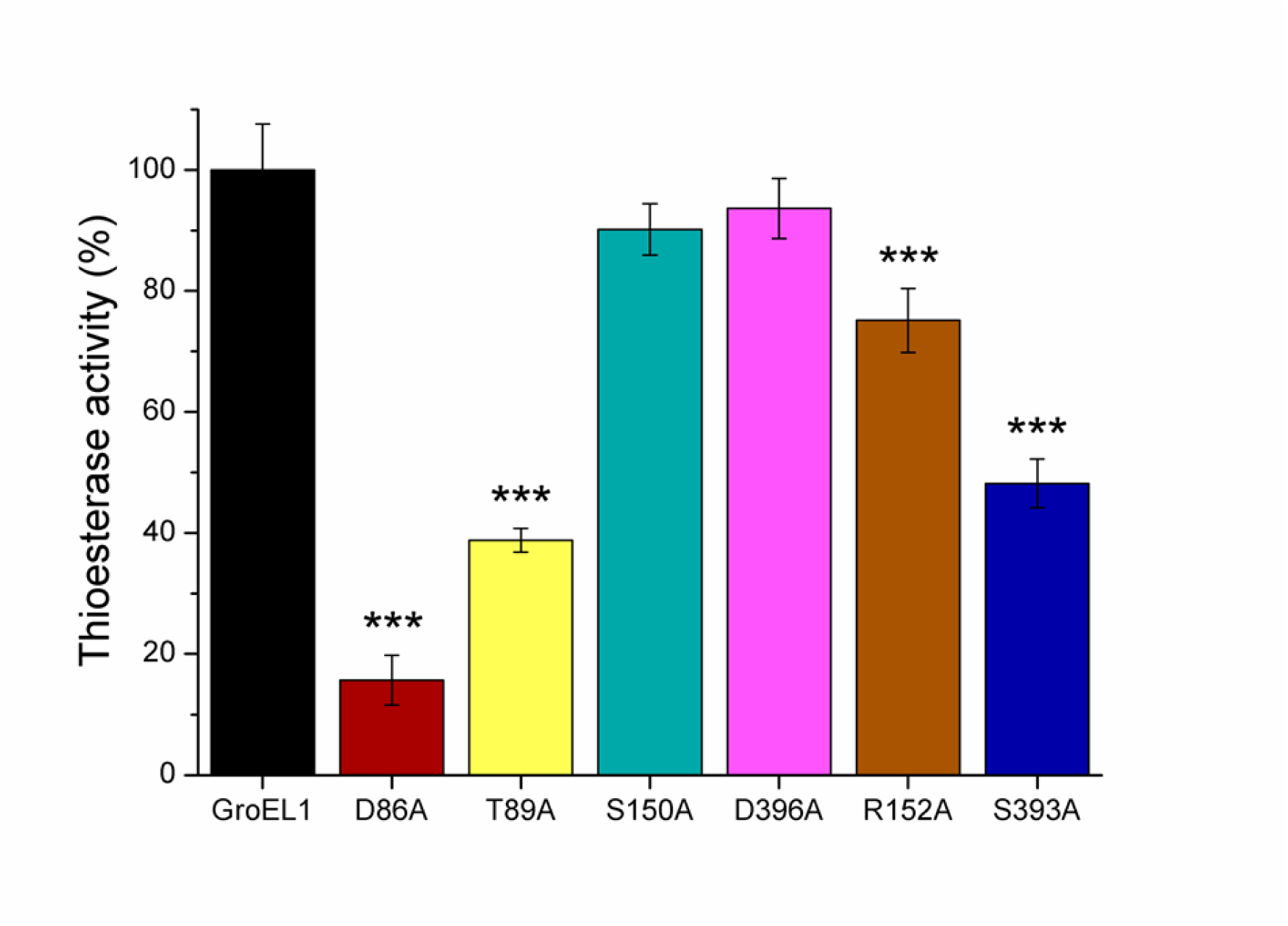
The thioesterase activity of GroEL1 mutant proteins in the presence of palmitoyl-CoA. The activity was monitored by measuring the absorbance at 412 nm for 1 h at 37 °C. The thioesterase activity of the D86A, T89A, S150A, D396A, R152A, S393A GroEL1 mutant proteins in comparison to the WT GroEL1 thioesterase activity. The mean from at least three independent experiments was calculated and plotted along with the standard deviation. The data were analyzed by two-sample unpaired *t* test: \*\*\**p* <0.001.

To elucidate the role of the C-terminal histidine-rich region (HDHHHGHAH) of GroEL1, in its thioesterase activity, we studied the activity of a recombinant *M. tuberculosis* GroEL1 devoid of its C-terminal histidine-rich region (GroEL1ΔHis). This mutant protein showed similar hydrolytic activities as the WT protein (Figure S9D). This region is thus not important for the hydrolytic activities of GroEL1, as previously reported for its ATPase activity^11^.

## Discussion

Here, we showed that GroEL/Hsp60 proteins are able of hydrolase activities, with low oligomeric forms of mycobacterial GroEL1 and Hsp60 being furthermore prone to use the long carbon chain substrate, palmitoyl-CoA. We also observed that the acyl chains released following these hydrolyses can eventually be transferred to the mycobacterial GroEL proteins, by auto-acylation.

Palmitoylation enhances protein hydrophobicity and can contribute to protein membrane insertion and as such can impact protein–protein interactions and protein-protein localization^49^. It will be interesting in the future to assess whether the Hsp60 membrane localization frequently observed in cancer cells could be due to auto-acyltransferase activity^50,51^. Although we could detect GroEL1 auto-palmitoylation, in the absence of pan-palmitoyllysine antibodies, we could not perform the classical PTM peptide enrichment strategy and were unable, using MS tools, to identify the palmitoylated residues. This could be due to the increased hydrophobicity of the PTM protein.

Considering that GroEL1 is required for PDIM biosynthesis in pathogenic mycobacteria^12,32^, that GroEL1 could eventually bind various PKS proteins, among others the PpsE protein involved in the last step of the phthiocerol biosynthesis required for PDIM biosynthesis and antibiotic resistance, we investigated whether GroEL1 could potentially facilitate the PpsE palmitoylation. Indeed, this could have an important impact on anchoring PpsE in the membrane. Palmitoylation analysis suggested that the combined presence of GroEL1 with the 60 kDa C-terminal region of the PpsE protein, containing various domains (PP, condensation, NRPS), improved the palmitoylation of the PpsE. This could partially explain the role of GroEL1 in PDIM biosynthesis and consequently also in vancomycin and rifampicin resistance^12^. Recent studies have highlighted the importance of proper assembling and translocation of lipids to the different layers of the mycobacterial cell wall^52^. As such, GroEL1 could play a crucial role in stress condition, by facilitating PpsE membrane anchorage through increased PpsE palmitoylation, reducing the risk of cytoplasm phthiocerol aggregation before transport through the cell membrane. GroEL/Hsp60 driven-PTM on native mycobacterial proteins, especially PpsE, should be further studied in the future.

We investigated possible binding modes of the adenylate moieties of the palmitoyl-CoA could fit in the ATP binding site of the GroEL1. The ATP-binding pocket seems conserved across GroEL proteins according to our MSA. Furthermore, we observed that ATP can inhibit the GroEL1 thioesterase activity and we hypothesized that it could be by competition. Looking to the structure of palmitoyl-CoA we hypothesized that this substrate of GroEL1 could even be twice hydrolyzed, once at the thioester bond, as observed in our thioesterase assay, but also between the two phosphates, when considering GroEL/Hsp60 ATPase or phosphodiesterase activity. This would eventually provide for three reaction products: adenosine 3’,5’-biphosphate, phosphopantetheine (that could also be used for polyketide synthase, among other PpsA-E proteins for building their operational acyl-carrier domain, ACP, required for PDIM biosynthesis) and the acyl-chain, used for PTM. This was studied in more detail based on docking calculations, carried out on a GroEL1 model, constructed using the *X. oryzae* GroEL structure, as template. Among the six residues selected because of potential involvement in GroEL1 enzymatic function, three residues, D86, T89 and S393, were identified as key players. Residues D86 and T89 are positioned in the main ATP-binding motif and may play a role in anchoring palmitoyl-CoA, which has an ATP-analog part. The mutation of these residues may have interrupted this favorable interaction, leading to thioesterase activity decrease. Further catalytic site prediction is limited because conformation changes after ATP or substrate interaction could take place, as previously observed for various thioesterase enzymes^53^. Oligomeric assembly regulation by ATP binding was thus both previously observed for GroEL proteins but also for some hotdog-fold thioesterases^53,54^. Indeed, in the *E. coli* GroEL–GroES but also in the human Hsp60-Hsp10 complexes, ATP binding induce tetradecamer complex conformational changes in order to switch from a substrate binding state to a substrate folding activity^55^. Interestingly, the S393, localized in the “central “ α-helix, intermediate domain, of the GroEL1 model, clearly involved in the GroEL1 thioesterase activity, was previously reported to be involved in conformational change through phosphorylation^19^. It is rarely present in other sequences, highlighting the uniqueness of GroEL1’s low oligomeric form. The fact that residue D396 is only involved in the activity of *E. coli* GroEL ATPase, not in ATP binding could explain why the mutation of this highly conserved residue did not influence GroEL1 thioesterase activity. For residues 150 and 152, no functional relevance could be inferred. These positions are less conserved, presenting eventually an A150 or a G152 in other GroEL proteins. Although this study focused on residues potentially involved in GroEL1 thioesterase catalytic sites according to GroEL1 model and docking prediction, further investigations are warranted, especially on position E461 as it could be involved in cooperativity between ATP binding and hydrolysis^56^. The residues potentially participating to GroEL protein oligomer stability (A2, E76, A109)^17,56–58^ could also be further investigated to assess whether they could provide an explanation for activity difference between GroEL1 and GroEL2 in *M. tuberculosis*.

Strikingly, several typical serine protease inhibitors (tetrahydrolipstatin, phenylmethylsulfonyl fluoride (PMSF), methoxy arachidonyl fluorophosphonate (MAFP)), similarly inhibited (around 50%) GroEL1 thioesterase activity as the S393A GroEL1 mutation (data not shown). This was also previously observed for *M. tuberculosis* TesA using serine protease inhibitors^39^. As half-of-site activity has been reported for various thioesterases^53^, it is tempting to hypothesize that the maximum 50% inhibition obtained by targeting a catalytic serine could be linked to a half-of-site reactivity. Inhibitors targeting other amino acid functions can be explored in the future.

It is worth mentioning that although D86 and T89 amino acid residues are probably in the ATP-binding pocket of GroEL1, GroEL1 has almost undetectable ATPase activity, which therefore impedes mutant ATPase activity analysis.

According to our study, GroEL1 could be a new type of thioesterase. Based on the catalytic mechanisms but also on the primary and tertiary structures of multiple thioesterases, various thioesterase classification attempts has been obtained^59^. The GroEL/Hsp60 chaperonin proteins do not fit into any of these families. However, as other thioesterases, we hypothesize that the various acyl-CoA substrates could first interact with some residues, in that case, the GroEL ATP-binding pocket (DGTTT), thereby initiating conformational change favoring the hydrolytic thioesterase activity.

This could have an important impact for anti-cancer or anti-tuberculosis drug developments. Indeed, various studies targeting *M. tuberculosis* GroEL1 and 2, aiming at treating tuberculosis, but also human mitochondrial Hsp60, to treat cancer and some autoimmune diseases^60,61^, were confronted to the cytotoxicity of their inhibitors, since they always targeted the classical, ubiquitous, folding or ATPase activity of those proteins. Instead, we propose to develop selective inhibitors of specific enzymatic activities only relevant for low oligomeric GroEL/Hsp60 (i.e. thioesterase activity on palmitoyl-CoA), without disrupting the ATPase activity. Although it is challenging to investigate the mechanism of low oligomeric GroEL/Hsp60 activity, we can hypothesize that the formation of the GroEL tetradecameric cage and steric hindrance could prevent palmitoyl-CoA to reach the thioesterase catalytic site of the high oligomeric forms.

## Conclusion

PTMs are reversible and modulate protein interactions, stability, localizations and activities. We show here for the first time that GroEL/Hsp60 proteins can drive such modifications due to their hydrolytic activities, including thioesterase and esterase activities, and their (auto)-acyltransferase activity. It will be important to assess whether these new GroEL/Hsp60 Swiss-knife activities could be essential tools to achieve their main goals, as chaperonins, in proper protein folding and localization and possible to target selective low oligomer GroEL/Hsp60-driven activities.

## MATERIAL AND METHODS

### *M. bovis* BCG strains and growth condition

Wild type *M. bovis* BCG (WT) and *M. bovis* BCG △cpn60.1 (or Cpn60.1/GroEL1 knockout, KO) strain precultures were performed in Middlebrook 7H9 medium (BD Difco) containing 10% (v/v) albumin-dextrose complex (ADC) and 0.05% ( v/v) Tween-80 (or 0.2% glycerol) at 37 °C^22^. The 7H9 precultures were used to launch potato (fresh uncooked French fries) precultures floating in 3.5%/6%(v/v), Sauton’s medium. Sauton’s medium contains 4.0 g asparagine, 2.0 g citric acid, 0.5 g K_2_HPO_4_, 0.5 g MgSO_4_, 0.05 g ferric ammonium citrate, 1.435 mg ZnSO_4_•7H_2_O, 60 mL/35 ml glycerol in 1 L, adjusted to pH 7.2 with 2 M NaOH. Colonies from the potato precultures were loaded on the interface air/liquid on 3.5%/6% Sauton’s medium and incubated at 37°C for 25 days for pellicle formation^38^.

### Plasmid constructions

*p*MtGroEL1, *p*MtGroEL1△His and *p*MtGroEL2 plasmids were previously obtained by cloning into the modified pET-15b vector^18^. This modified vector includes a cleavage site for the rhinovirus 3C protease after the 5×His-tag sequence^18^. Coding sequences for the *E. coli* GroEL and the human Hsp60 were amplified by polymerase chain reaction (PCR) and inserted in the modified pET-15b vector using the restriction sites *Nde*I and/or *Xho*I to obtain the *p*MtGroEL and *p*MtHsp60 plasmids, respectively. The primers used are shown in supplementary data (Table S1). All the plasmid constructions were verified by sequencing.

### Obtention of *p*MtGroEL1 mutated plasmids

The oligonucleotide primers designed to insert individual mutations in the *cpn*60.1 sequence using QuikChange Lightning Site-Directed Mutagenesis Kit (Agilent) are shown in supplementary data (Table S2). Mutagenized *p*MtGroEL1 plasmid sequences were amplified by PCR, before *Dpn* I digestion to remove parental plasmid DNA. After digestions and bacterial transformation, plasmids were sequenced to identify *p*MtGroEL1-D86A, *p*MtGroEL1-T88A, *p*MtGroEL1-S150A, *p*MtGroEL1-R152A, *p*MtGroEL1-S393A and *p*MtGroEL1-D396A transformed colonies.

### *pMt*PpsE plasmid construction

Two carboxyl terminus PpsE sequences, encoding either the phosphopantetheine (PP)-binding domain combined with the condensation domain or the condensation and the non-ribosomal peptide synthetase (NRPS) domains, were cloned by PCR using primer PP, COND, NRPS (Table S3) in the modified pET-15b vector. This allowed to obtain the p*Mt*PpsE(PP-COND) and the p*Mt*PpsE(COND-NRPS) plasmids, respectively. As the p*Mt*PpsE(COND-NRPS) was devoid of the 129 bp encoding for the last 42 amino acid residues, an additional plasmid was obtained (p*Mt*PpsE-end) by PCR, by cloning the last 129 bp of the ppsE gene sequence into the p*Mt*PpsE(COND-NRPS), using primer END (Table S3). The p*Mt*PpsE-end plasmid allows to produce the C-terminal end of the PpsE protein with the condensation and NRPS domains. Finally, a new plasmid was constructed by insertion of the PpsE C-terminus region (about 900 bp) into the p*Mt*PpsE(COND-NRPS), allowing to obtain the p*Mt*PpsE-C plasmid used to produce the last 555 amino acid residues of PpsE. The PpsE gene sequences were verified by sequencing.

### Production and purification of recombinant proteins

The *E. coli* strain BL21(DE3) transformed with either the p*Mt*GroEL, the wild-type or mutated *p*MtGroEL1, the *p*MtGroEL2 or the *p*MtHsp60 plasmid were used for overproduction of recombinant proteins. The bacteria were grown in Lennox broth (LB) medium (Carl Roth Gmbh, Karlsruhe, Germany) with or without 100 μg/mL ampicillin. Gene expression was induced by addition of 1 mM isopropyl β-D-1-thiogalac-topyranoside (IPTG) when the optical density of the bacteria cultures reached 0.4-0.6 at 600 nm. Bacteria cultures were further incubated 20 h at 18°C for GroEL1 production or at 30°C for GroEL2 and Hsp60 production, or at 37°C for GroEL production. Bacteria were harvested by centrifugation at 4000 g at 4°C and resuspended in a lysis buffer (20 mM HEPES, 300 mM NaCl, 10 mM imidazole, pH 8.0, EDTA-free protease inhibitor cocktail from Carl Roth Gmbh, 20 mM MgSO_4_ and 10 μg/mL DNase from Sigma). The suspension was homogenized in a Potter-Elvehjem homogenizer before lysis of the bacteria by three passages through an Emulsiflex-C3 (Avestin). The bacterial lysates were centrifuged 40 min at 30,000 g at 4°C. The supernatant was loaded on a 1 mL Ni-NTA agarose column (Thermo Scientific) and incubated under gentle agitation during 1 h at 4°C. The column was washed for the first time with 20 mM HEPES, 300 mM NaCl and 10 mM imidazole (pH 8.0), for the second wash with 20 mM HEPES, 300 mM NaCl and 40 mM imidazole (pH 8.0), before protein elution with 20 mM HEPES, 300 mM NaCl and 250 mM imidazole (pH 8.0). Imidazole was removed from the protein solutions by exchange with 20 mM HEPES, 150 mM NaCl (pH 7.5) using PD-10 desalting columns (Cytiva). To remove the His-tag of the proteins, HRV-3C protease (protein/enzyme mass ratio of 75:1) was added for a 4°C overnight incubation. Proteins were further purified by size exclusion chromatography (SEC) using a Superdex 200 10/300 GL analytical column connected to an ÄKTA purifier system (Cytiva). The protein concentrations were determined by measuring the absorbance at 280 nm using the calculated extinction coefficients of 16,960 M^-1^ cm^-1^ for GroEL1, 15,930 M^-1^ cm^-1^ for GroEL2, 10,430 M^-1^ cm^-1^ for GroEL and 15,930 M^-1^ cm^-1^ for Hsp60. The recombinant proteins were verified by SDS-PAGE and mass spectrometry for purity and integrity, respectively (for details see open access data on Zenodo DOI : 10.5281/zenodo.13355298). Recombinant proteins were flash-frozen in liquid nitrogen and stored at -80 °C for further experiments.

### Native-PAGE (Native Polyacrylamide Gel Electrophoresis)

Native-PAGE was performed at a constant current of 20 mA/gel using Mini-PROTEAN^®^ TGX™ precast 4-15 % gels (Bio-Rad, USA) under native conditions. Molecular weight estimation of proteins separated on the native-polyacrylamide gels was realized using NativeMark^TM^ Unstained Protein Standard (Invitrogen).

### ATPase activity assay

The ATPase activity of the purified recombinant *M. tuberculosis* GroEL1 and *E. coli* GroEL were quantified by colorimetry as previously described, with minor modification^62^. Briefly, 10 μM proteins were incubated 1 hour at 37 °C with 100 μL reaction buffer containing 10 mM KCl, 2 mM ATP, 10 mM MgCl_2_, 20 mM HEPES (pH 7.5) and 150 mM NaCl. Enzymatic reactions were terminated by adding 500 μL 1% SDS. Two hundred μL of ammonium molybdate reagent were first added to each reaction immediately followed by the addition of 200 μL Elon reagent. After 15 min incubation at room temperature, absorbances were measured at 700 nm (Tecan, Männedorf, Switzerland). Controls without proteins were always included.

### Thioesterase activity assays

Thioesterase activity assays were performed in 96-well plates and the thiol group enzymatic production was highlighted using the 5,5’-dithiobis(2-nitrobenzoic) acid (DTNB) able to yield a DTNB-derived yellow product^63,64^. Acetyl-CoA, propionyl-CoA, decanoyl-CoA and palmitoyl-CoA were used as substrates. The reaction medium contained 100 mM sodium phosphate (pH 7.2), 2 mM DTNB, 10 μM proteins and increasing substrate concentrations in a final volume of 200 μL. The thioesterase activity was monitored spectrophotometrically (Tecan) over time at 412 nm. The activity was described in international units (U), corresponding to 1 μmol/min released TNB^2-^. Here, it was first calculated for one mL (unit/mL) taking the dilution factor (df) at each time point in min. (t), the extinction factor ε_412_=14,150 M^-1^ cm^-1^, the absorbance (A) at 412 nm, the used volume of enzyme (V_e_ in mL) and the volume of reaction (0.2 mL) in consideration, using the formula:

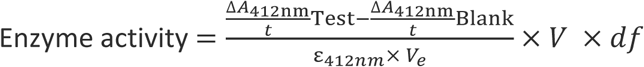

Then, the specific activity (SA) was calculated in unit/mg enzyme, knowing the concentration of the enzyme (mg/mL). Curve fitting and evaluation of apparent K_m_ and apparent V ^65^ were done using OriginPro 8.0 enzyme kinetics program^39^.

### Esterase activity assays

The esterase activity assays were performed according to Nguyen *et al*.^66^. Enzymatic reactions were carried out over a period of 60 min in a 96-well plate in 100 mM sodium phosphate (pH 7.6) with 10 μM proteins and various substrate concentrations. The 4-nitrophenyl acetate (4-NPA), 4-nitrophenyl butyrate (4-NPB) and 4-nitrophenyl palmitate (4-NPP) were used as substrates. The absorbance was measured at 410 nm (spectrophotometer, Bio-Tex) every 30 s for 60 min at 37°C.

### Thermal stability assay

The SYPRO orange dye (Invitrogen) was used to monitor protein unfolding during heating. The thermal shift assay was conducted in 96-well plate using the CFX96^TM^ real-time system (Bio-Rad, USA) under different conditions. Five μM GroEL1 and SYPRO orange dye (2000 times dilution of the commercial stock) were present in all reactions in the presence of 150 mM NaCl and 20 mM HEPES buffer (pH 7.5). The samples were heated from 10 °C to 95 °C with a heating rate of 1°C/min. The fluorescence intensity was recorded at Ex/Em = 465/510 nm. The data were obtained from the Bio-Rad Precision Melt Analysis software 1.0. Plots of d*F*/d*T* (fluorescence intensity difference, d*F*, in function of temperature difference, d*T*) were obtained using the OriginPro 8 software.

### Structural modelling of GroEL1

To perform docking calculation several models of GroEL1 structures were generated by comparative modelling with Modeller 9v4^67^. First a BLAST search performed on the Protein Data Bank (PDB) sequences identified several proteins as potential templates, notably GroEL2 from *M. tuberculosis* (PDB ID: 3rtk) and GroEL from *Xanthomonas oryzae* (PDB ID: 6kvf) and from *E. coli*, the latter in complex either with an ATP analogue (PDB ID: 1sx3) or with GroES (PDB ID: 1pcq). Sequence alignments were carried out with ClustalW^68^. A model built by AlphaFold2^69^ was also used.

### Docking calculations

The structures were prepared with the Protein Preparation Wizard workflow implemented in the Schrödinger package^70^. The initial 3D structures of the ligands were generated using the Ligprep module (Schrödinger, LLC, New York, NY, USA, 2018).

In the present work, the binding region in the monomer was defined by a box centered on a pocket delimited after superposition of a GroEL *E. coli* X-ray structure determined with phosphothiophosphoric acid-adenylate ester (AGS), an ATP analog (PDB ID: 1sx3) on the modelled structure. Docking was performed using the Glide XP docking protocol and scoring function which approximates a systematic search of positions, orientations, and conformations of the ligand in the receptor binding site using a series of hierarchical filters. The default settings of Glide were used.

### Phylogenetic tree building

53.000 sequences of chaperonins GroEL and TCP1 were retrieved from Uniprot^71^. This dataset was filtered first for the most scientifically relevant bacterial and archaea genera, as well as

Eumetazoa. A sequence-based data reduction was performed using the software CD-Hit^72^ with an 80% sequence similarity criterion. The resulting dataset encompassed 698 sequences. These sequences were aligned using the novel pipeline SIMSApiper^73^, which uses protein structure models to overcome the high sequence divergence in the dataset. IQ-Tree2^74^ was used to calculate an unrooted phylogenetic tree from this MSA with ModelFinder^75^ and and 10.000 ultra-fast bootstrap replicates^76^. The root was placed between the cluster of archaeal and bacterial sequences. The resulting tree was visualized using TreeViewer^77^.

### Contextualization of GroEL1 mutations

Residues with functional implications on GroEL function in *E. coli* (P0A6F5) and *M. tuberculosis* (P9WPE9) were collected from Uniprot. These residues were mapped onto the multiple sequence alignment (MSA) and visualized as sequence logos^78^. Each sequence logo provides information of the consensus sequence at the position, allowing to better visualize sequence conservation across GroEL/Hsp60 and TCP-1 proteins and to infer possible functional implications of these residues.

### Recombinant GroEL1 auto-acylation characterization

The potential acylation of recombinant GroEL1 was investigated by mass spectrometry. Briefly, 10 μM GroEL1 in esterase reaction or 89 μM GroEL1 in thioesterase reaction were incubated with 2 mM 4-NPA or 4-NPB substrate or with 1 mM propionyl-CoA in 100 mM sodium phosphate pH 7.6 and the reaction mixture was incubated at 37°C for 1 hour.

After the esterase enzymatic reaction, the buffer was exchanged for 75 mM NaCl and 10 mM HEPES pH 7.5 using a Vivaspin 6 ultrafiltration spin column (10 000 MWCO). The intact mass of the proteins was determined by mass spectrometry. After digestion with trypsin/Lys-C or EndoGlu-C endoproteases (sequencing grade, Promega) in an enzyme/protein mass ratio of 50:1, the modified peptides were analyzed by tandem mass spectrometry (MS/MS).

After the thioesterase reaction, an additional step for product enrichment was performed by immunoprecipitation directly.

### Peptide immunoprecipitation with antibody conjugated agarose beads

The digestion of native or recombinant GroEL1 proteins was carried out by addition of trypsin. First, 1 mg dry tryptic peptides was dissolved in 100 μl IP buffer (100 mM NaCl, 1 mM EDTA, 20 mM Tris-HCl, 0.5% NP-40, pH 8.0) and insoluble proteins were discarded by centrifugation at 12,000×g for 10 min at 4°C. The supernatant was then incubated overnight under gentle rotation at 4°C with 20 μL of anti-propionyllysine antibody conjugated agarose beads (PTM BIO), previously washed with 0.5 mL of ice-cold PBS. Afterwards, antibody conjugated beads were collected by centrifugation at 500×g for 30 s and washed first by adding 0.5 mL wash buffer I (100 mM NaCl, 1 mM EDTA, 20 mM Tris-HCl, 0.5%, v/v, NP-40, pH 8.0), followed by adding 0.5 mL wash buffer Ⅱ (100 mM NaCl, 1 mM EDTA, 20 mM Tris-HCl, pH 8.0) and finally using 0.5 mL Milli-Q water. The bound peptides were eluted with 100 μl of elution buffer (0.1%, v/v, trifluoroacetic acid) and dried for mass spectrometry analysis.

### Analysis of GroEL1 protein PTM by LC-MS/MS

After reduction and alkylation, GroEL1 was digested with trypsin at a ratio of 1:25 (enzyme: substrate) and incubated overnight at 37°C. Trypsin digest was then stopped with 1ml 5% (v/v) formic acid. Tryptic peptides were analyzed on an ultra-high-performance liquid chromatography–high-resolution tandem mass spectroscopy (UHPLC-HRMS/MS) system (Eksigent nanoLC 425 - AB SCIEX TripleTOF™ 6600+). Tryptic peptides (2 µl) were separated in a 15 cm C18 column (Triart C18, 3µm, YMC) using a linear acetonitrile (ACN) gradient [5-35% (v/v), in 75 min] in water containing 0.1% (v/v) formic acid at a flow rate of 5 µl/min.

Mass spectra (MS) were acquired over the range of 400-2,000 m/z in high-sensitivity mode (resolution > 30,000), with a 250 msec accumulation time. The instrument was operated in DDA (data dependent acquisition) mode, and MS/MS spectra were acquired over the range of 100-2,000 m/z. The precursor selection parameters were as follows: intensity threshold, 100 cps; maximum precursors per cycle, 30; accumulation time, 50 msec; and exclusion time after two spectra, 4 sec. These parameters lead to a duty cycle of 1.7 sec per cycle to ensure that high quality extracted ion chromatograms (XICs) were obtained for peptide characterization. ProteinPilot Software (v5.0.1 – ABSciex, United States) was used to perform database searches against the GroEL1 sequence. The search parameters included propionylation pre-digestion as special factor, all biological modifications, missed trypsin cleavage sites. All identification with a coverage of 99% (95% confidence), a mass accurate score below than 20 ppm and a clear MS/MS spectrum were considered.

### GroEL1 and PpsE-C (auto-)palmitoylation characterization

Thioesterase reactions (20 μL) containing 100 mM sodium phosphate (pH 7.2), 10 μM recombinant mycobacterial proteins and 100 μM fluorescent related palmitoyl-CoA, the 16-NBD-16:0, (N-[(7-nitro-2-1,3-benzoxadiazol-4-yl)-methyl]amino) palmitoyl Coenzyme A (Avanti), were incubated at 37°C for 1 h. Afterwards, samples were diluted and inactivated for 4-15 % native-PAGE followed by fluorescence analysis (ChemiDoc XRS+, Bio-Rad). The most intensive fluorescent bands were cut from the gels and put in trypsin digestion reactions as described above. The tryptic peptides were analyzed by LC-MSMS, as described above. The relative abundance of GroEL1 or PpsE were analyzed using the ProteinPilot software based on the intensity of identified peptides.

### Production and purification of native *M. bovis* BCG GroEL1

*M. bovis* BCG biofilm cultures were grown on zinc deficient Sauton’s medium containing 6% (v/v) glycerol to obtain large amounts of secreted proteins^79^. The culture medium was clarified by filtration through a filter unit of 0.22 μm porosity (Millipore) and concentrated using a Pellicon filter unit equipped with membranes of 30 kDa cut off. To obtain the native GroEL1 having a natural His rich C-terminal region, the retained compounds from the Pellicon filtration were loaded on a Ni-NTA agarose column (Thermo Scientific) for further purification as described above.

### Fourier transform infrared spectroscopy (FTIR)

ATR_FTIR spectra were recorded on a Bruker Equinox 55 spectrophotometer equipped with a MCT detector at a resolution of 2 cm^-1^. The spectra were obtained in the ATR mode by using a Golden Gate™ ATR accessory (Specac, Orpington, United Kingdom) with an integrated total reflection element composed of a single diamond. The angle of incidence was 45°. The spectrophotometer was continuously purged with dry air and measurements were carried out at 21°C. Thin protein films were obtained by slowly evaporating samples under a stream of nitrogen. For hydrogen/deuterium exchange, the sample was flushed with D_2_O saturated N_2_. We recorded 64 scans every minute for 2 h and averaged for each time point for kinetic measurements. Data were processed using the Kinetics program developed by Dr. Erik Goormaghtigh^80^ in MatLab R2007a.

### Statistical Analysis

Figures were prepared using OriginPro 8. Unpaired t-test statistical analysis was performed using OriginPro 8 software unless otherwise stated. A *p* value < 0.05 was considered statistically significant. Experimental results are shown from pooled data of at least three independent experiments.

## Acknowledgments

We thank Stephane Canaan (CNRS, LISM, IMM FR3479, Aix-Marseille University, France), and Didier Vertommen (de Duve Institute and MASSPROT platform, UCLouvain, Brussels, Belgium) for critical reading.

The PhD study of Z. Zhou and D. Yang were partially supported by the China Scholarship Council (CSC) (No. 201908210292), the Van Buuren Prize, and the De Meurs-Francois Prize. Isaline Lambert was supported by the “Amis des Instituts Pasteur à Bruxelles” association and the F.R.S.-FNRS. The Bioprofiling platform used for proteomic analysis was supported by the European Regional Development Fund and the Walloon Region, Belgium.

## Author contributions

VF and GV conceived and supervised the research; VF and DY developed initial concepts; ZZY, DY, VF and GV designed the experiments; ZZY, GY, DY, CD, CM, MP and RW performed the research; ZZY, DY, GV, IL, MP, RW, SLH and VF analyzed the data; ZZY and VF drafted the manuscript; ZZY, GV, YD, JFF, SLH, MP and VF revised the manuscript; RW, GV and VF provided materials.

## Declaration of interests

The authors declare no competing interests.

**Source data** available on Zenodo, DOI: 10.5281/zenodo.13355298

---

**Figure 1.**
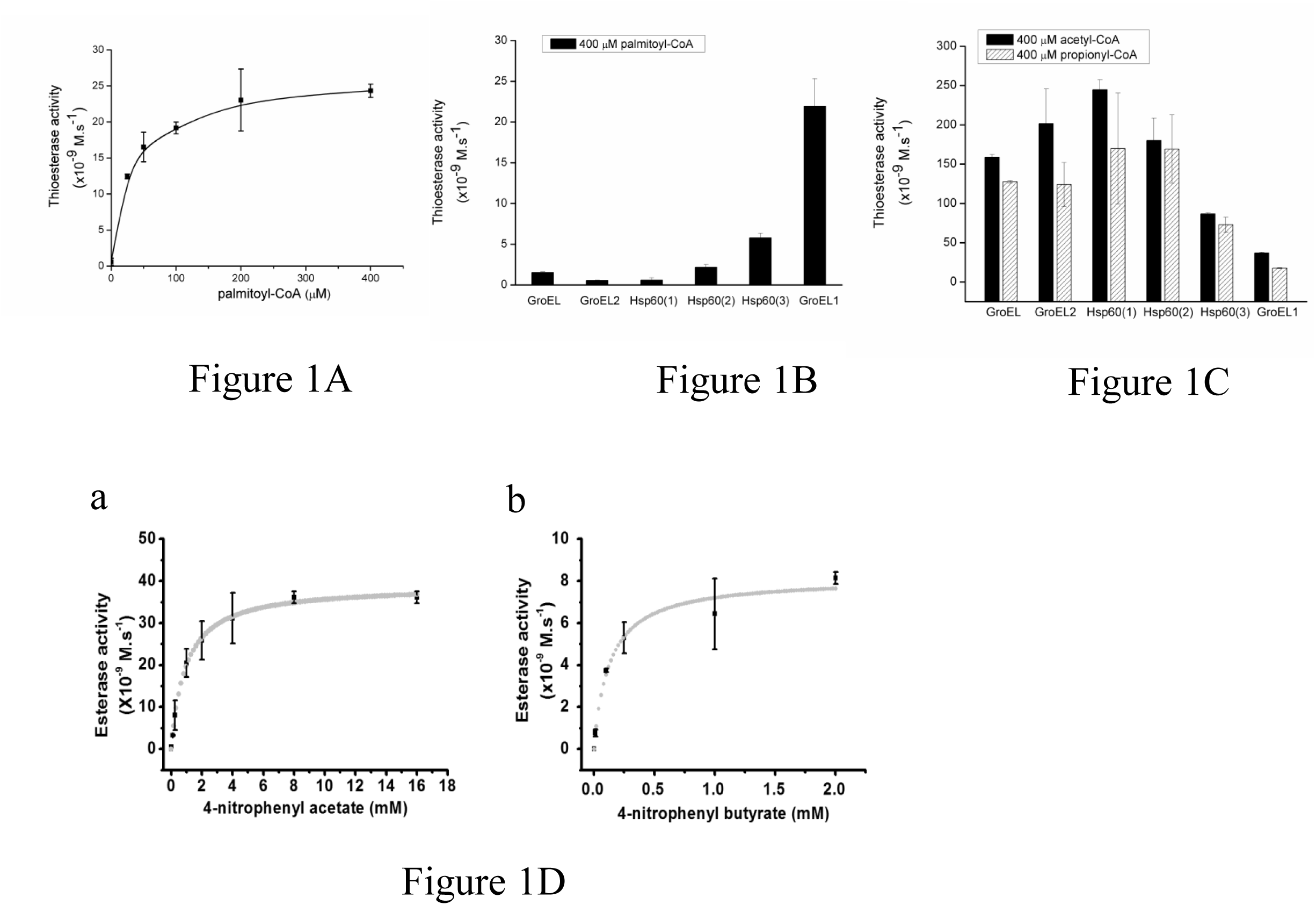
Thioesterase/esterase activities of recombinant proteins. (A) Dose-dependent thioesterase activity of recombinant GroEL1 in the presence of palmitoyl-CoA, (B) Comparison of different recombinant chaperonin thioesterase activity in the presence of 400 μM palmitoyl-CoA, (C) Recombinant chaperonin thioesterase activity in the presence of 400 μM acetyl-CoA and propionyl-CoA. (D) GroEL1 esterase activity. in the presence of 4-nitrophenyl acetate (a) or 4-nitrophenyl butyrate (b). The data were obtained from at least three independent experiments.

**Table 1.**
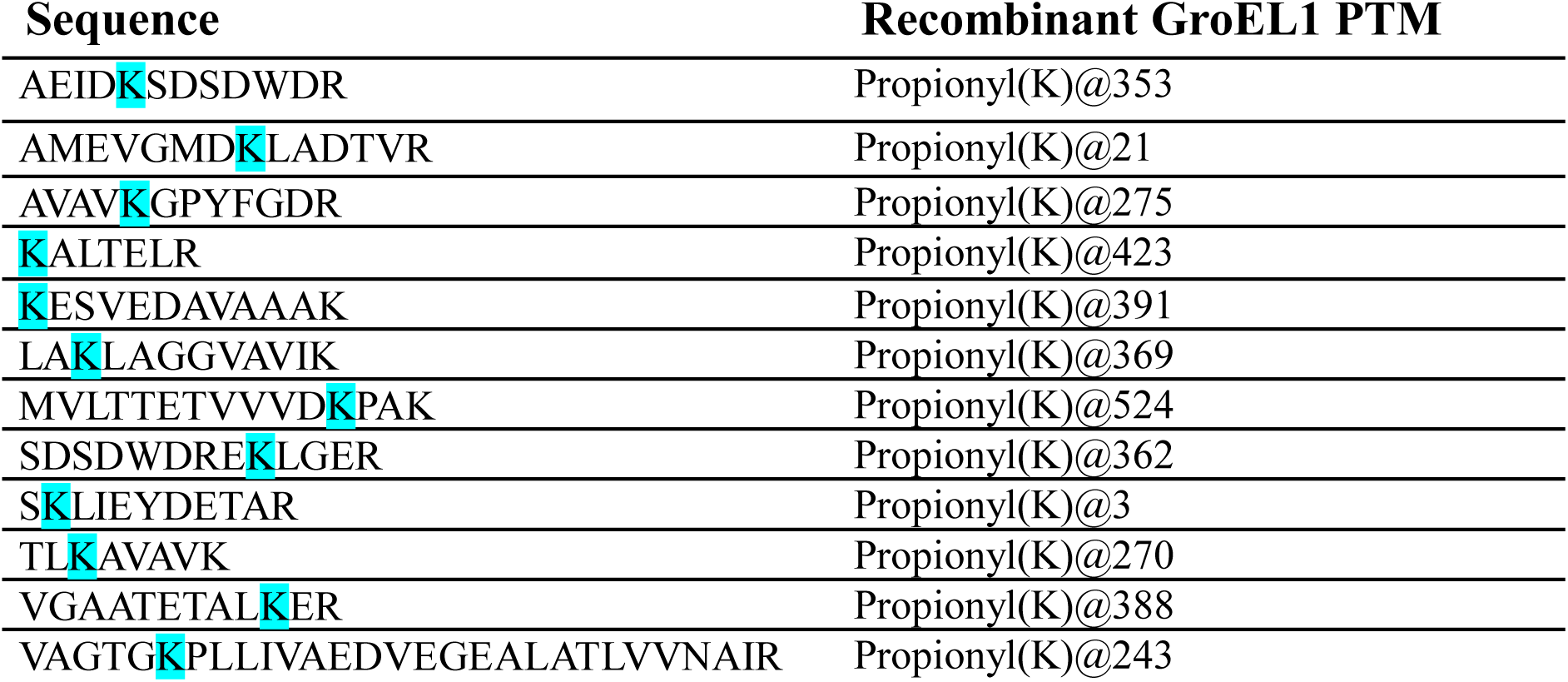
Propionylations on recombinant GroEL1 protein.

**Table 2.**
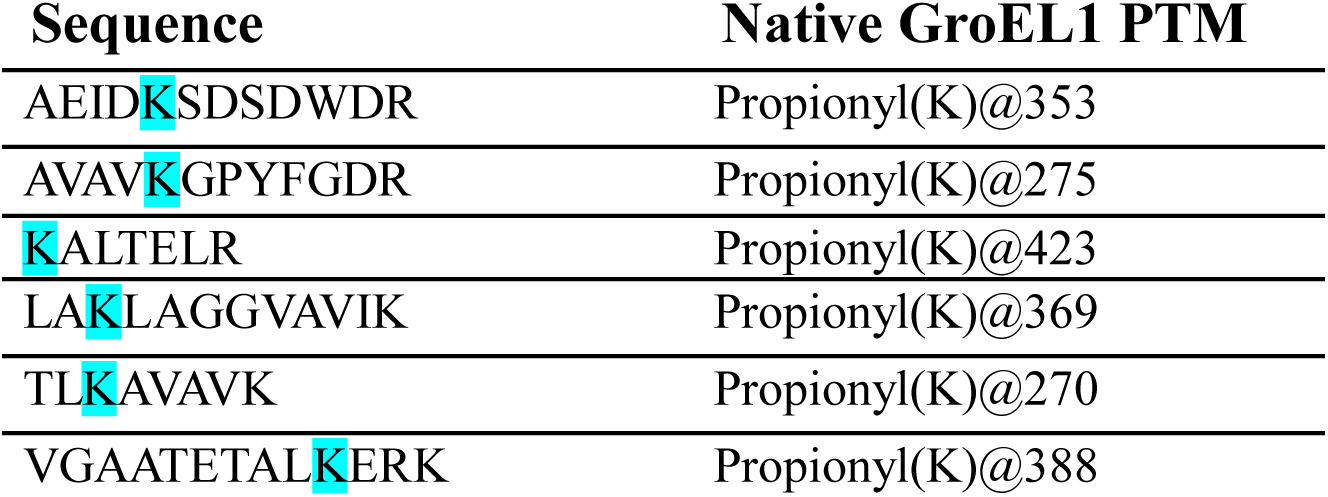
Propionylations on native *M. bovis* BCG GroEL1.

**Figure 2.**
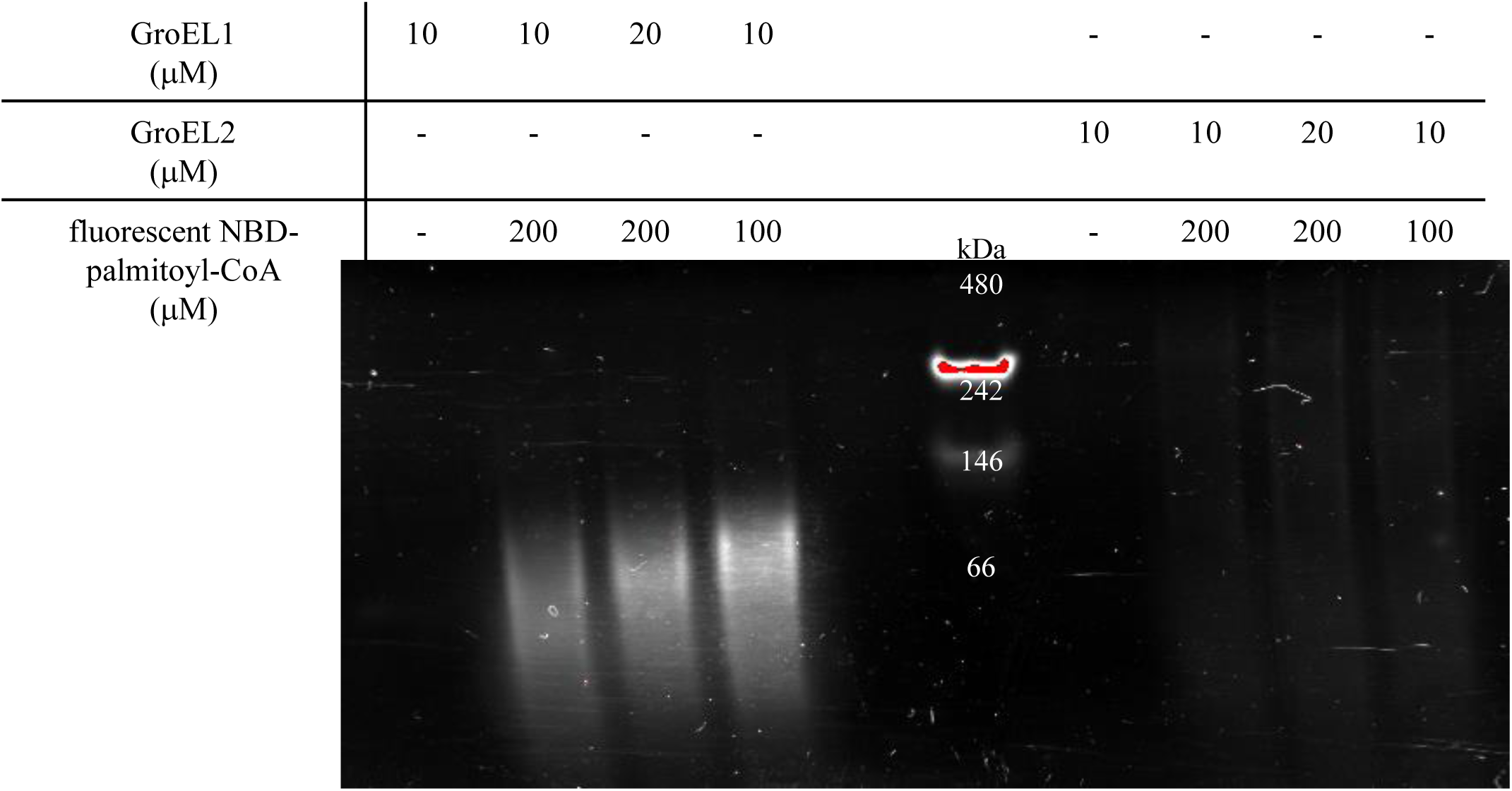
Detection by fluorescence of auto-palmitoylation of recombinant mycobacterial GroEL proteins. The reactions containing 100 mM sodium phosphate (pH 7.2), 10 μM recombinant mycobacterial proteins and 100 μM fluorescent related palmitoyl-CoA, incubated at 37°C for 1 h, were analyzed by 4-15 % native-PAGE followed by fluorescence analysis. The data are representative of at least three independent experiments.

**Figure 3.**
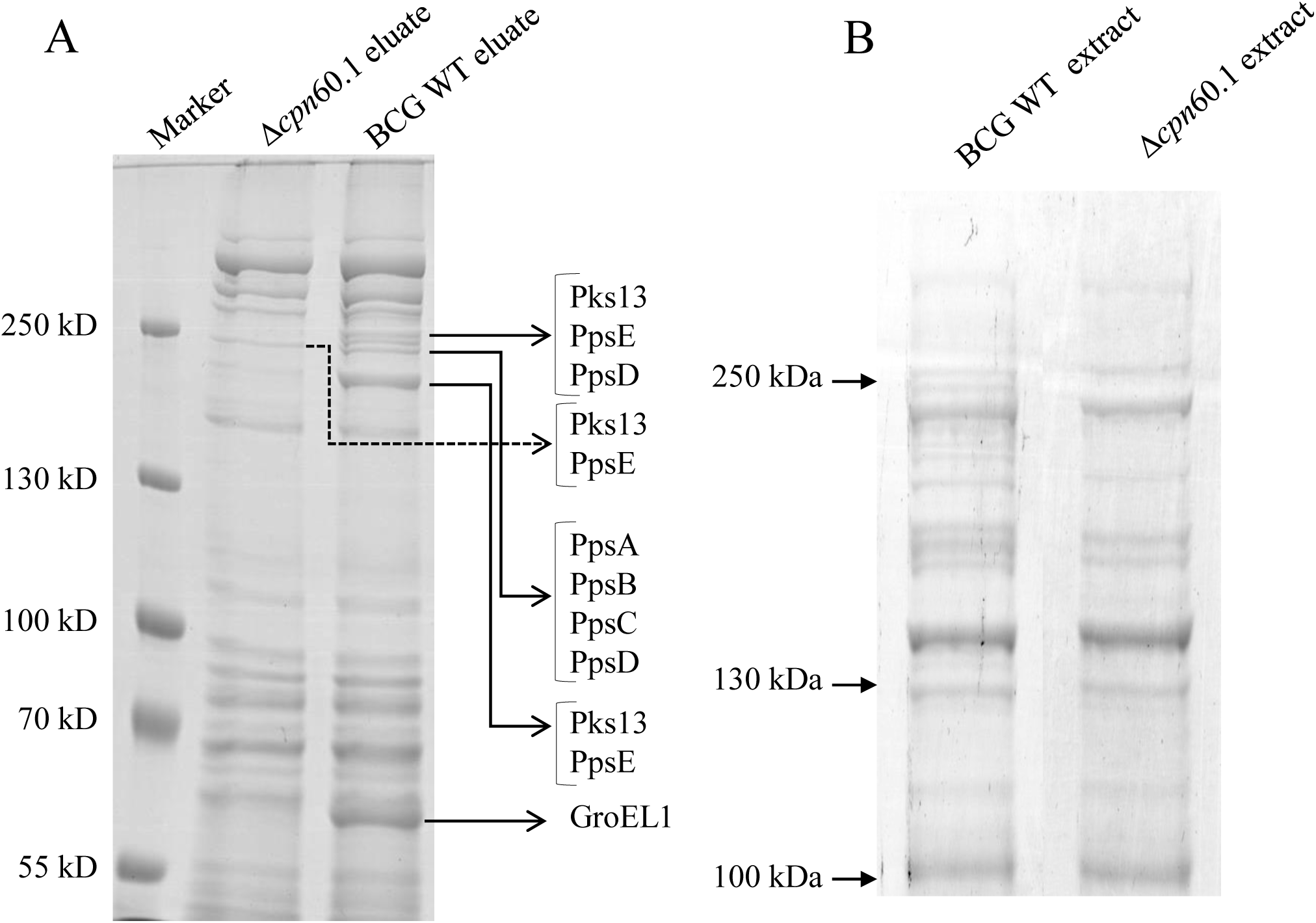
Co-elution of GroEL1 and PKS proteins from the WT and Δ*cpn*60.1 *M. bovis* BCG. (A) Protein eluates after Ni-column purification were analyzed by 12 % SDS-PAGE, and further characterized by MS/MS after gel elution and trypsin digestion. (B) Protein extracted from the WT and Δ*cpn*60.1 *M. bovis* BCG according to 12 % SDS-PAGE analysis.

**Figure 4.**
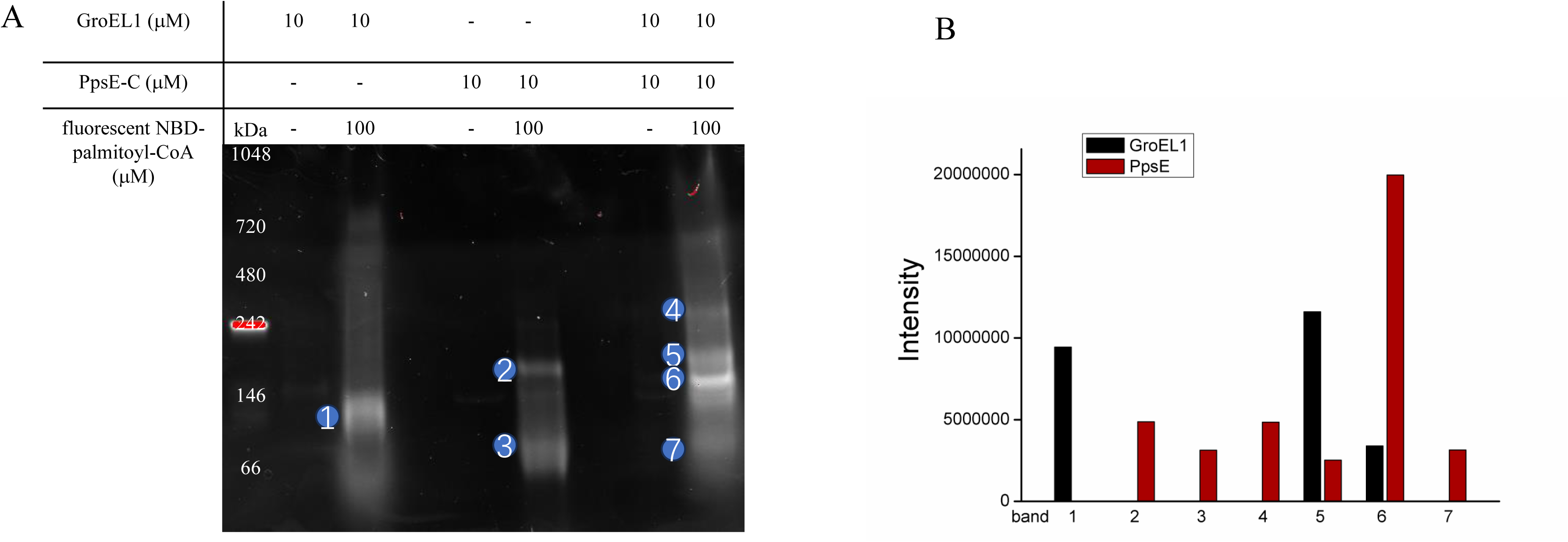
Palmitoylation of recombinant GroEL1 and PpsE-C proteins. (A) Fluorescence detection of palmitoylated proteins on 4-15 % native-polyacrylamide gel after 1h incubation at 37 °C in 100 mM sodium phosphate (pH 7.2) in the presence of eventually 10 μM recombinant GroEL1 and/or 10 μM PpsE-C protein and 100 μM fluorescent related palmitoyl-CoA. (B) LC/MSMS identification analysis of fluorescent digested proteins after elution from the native polyacrylamide gel (A).

**Figure 5.**
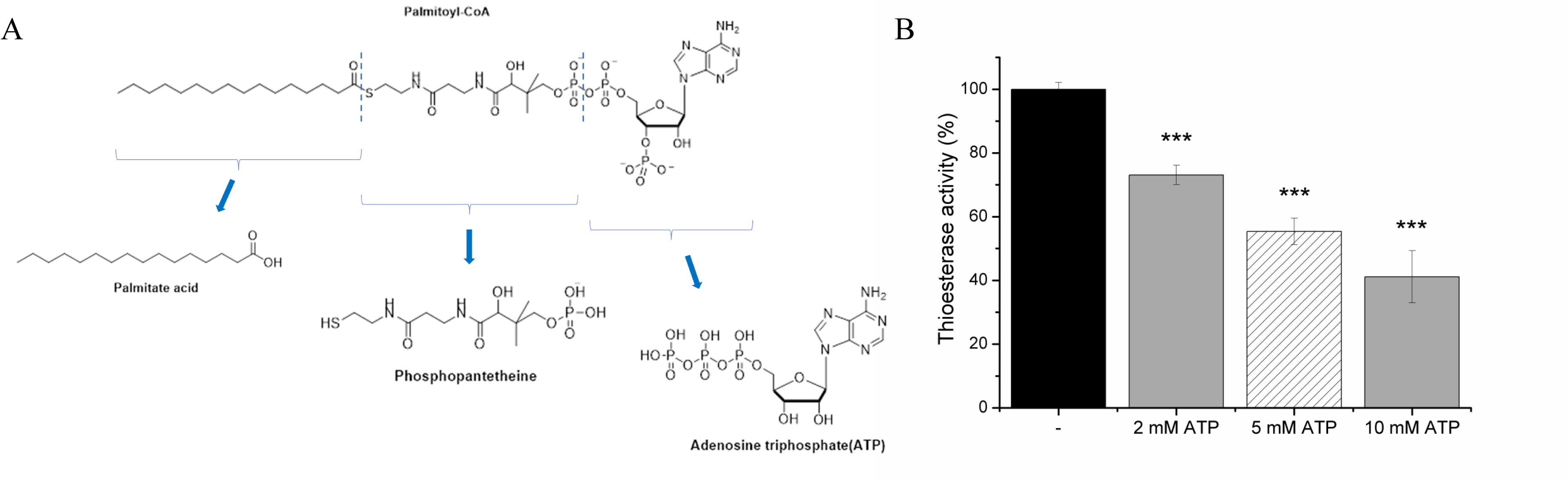
(A) Palmitoyl-CoA structure. (B) Dose-dependent impact of ATP on GroEL1 thioesterase activity. The activity was monitored by measuring the absorbance at 412 nm for 30 min at 37 °C. The data were obtained from at least three independent experiments. \*\*\**p*<0.001.

**Figure 6.**
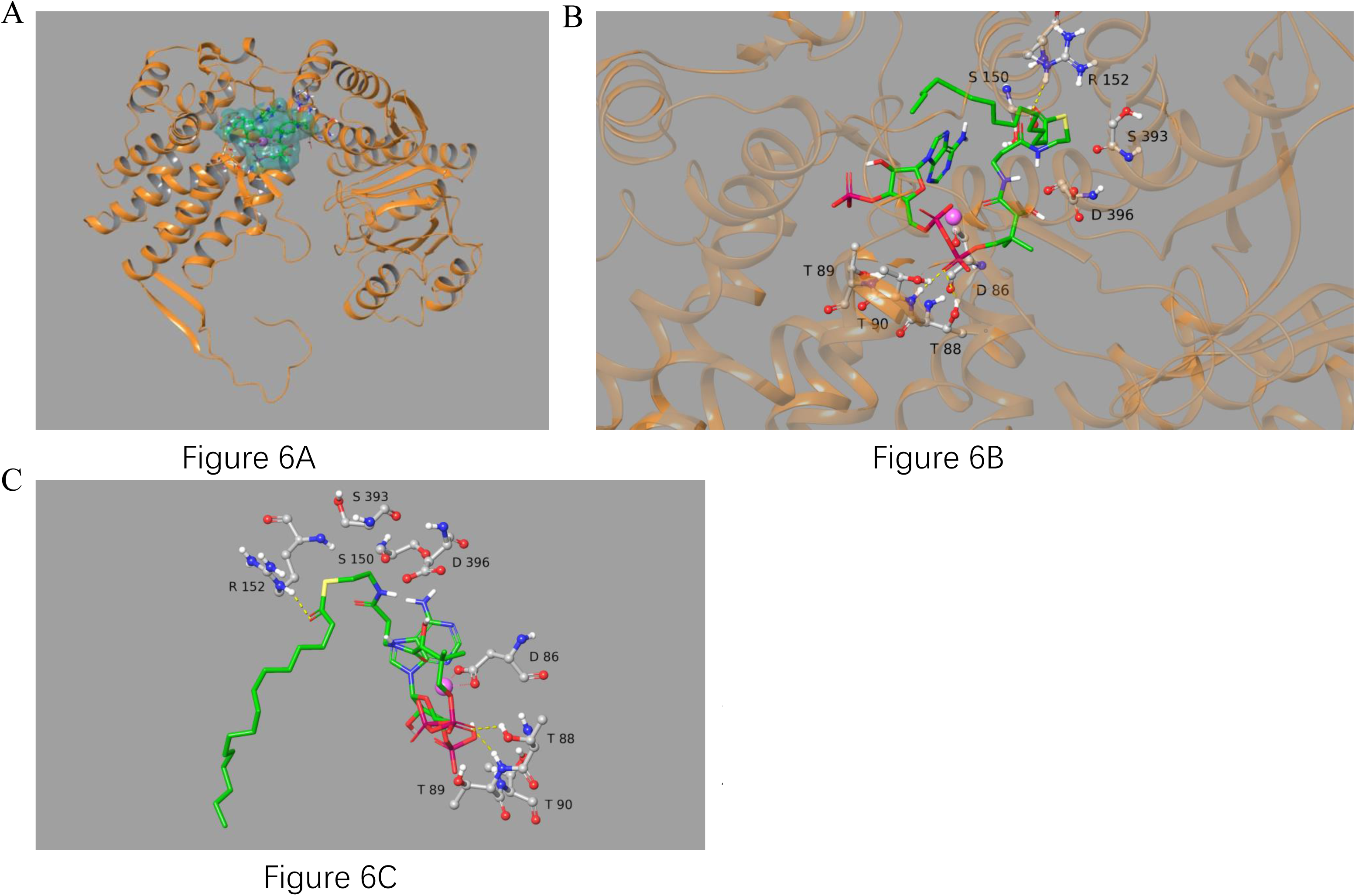
(A) Palmitoyl-CoA was docked in the ATP binding pocket of the model from *Xanthomonas oryzae* in the presence of magnesium. (B,C) The positions predicted in the structure modeled with GroEL *Xanthomonas oryzae* in the presence of magnesium were shown.

**Table 3.**
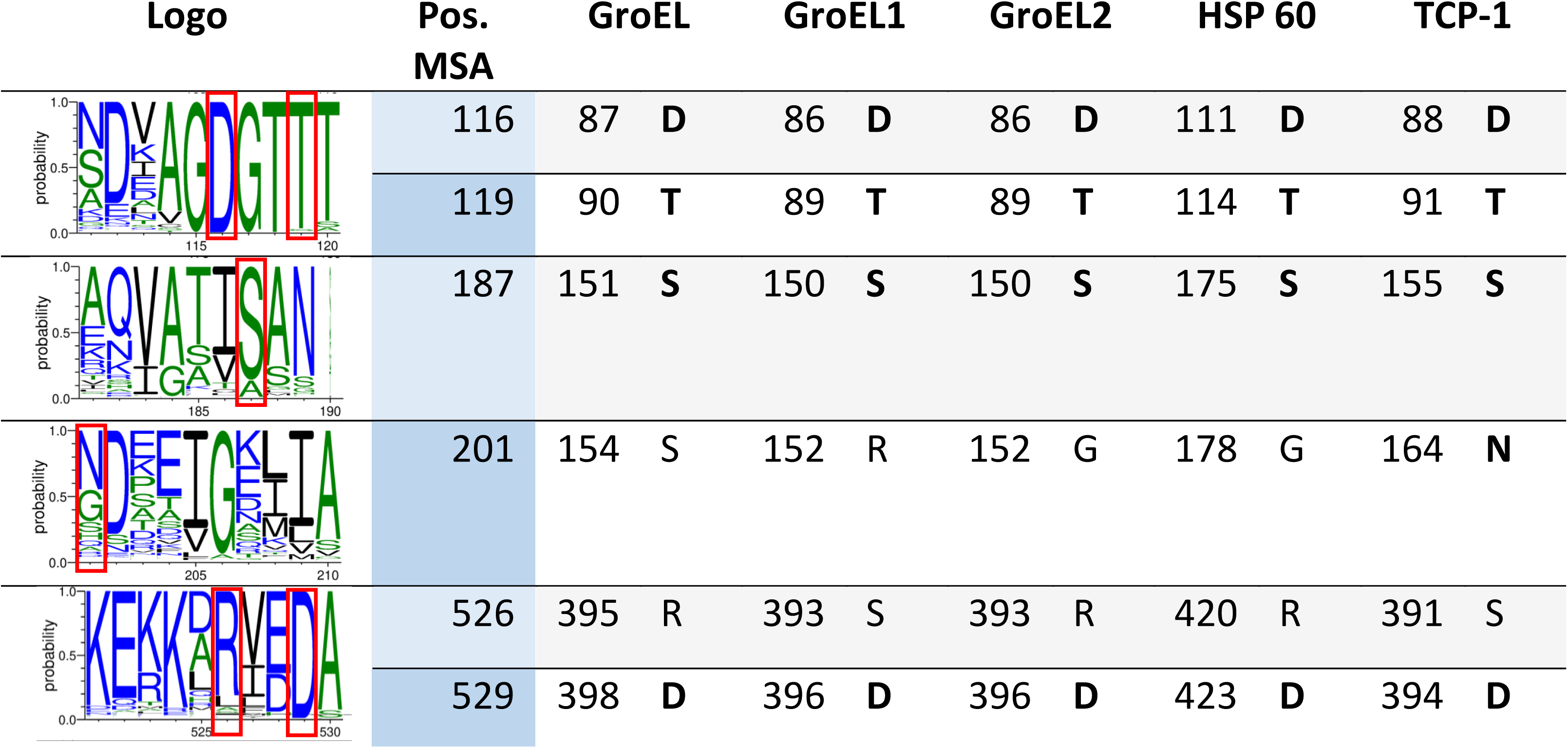
Residues mutated in GroEL1 mapped via the global MSA onto corresponding residues in other proteins of interest.

**Figure 7.**
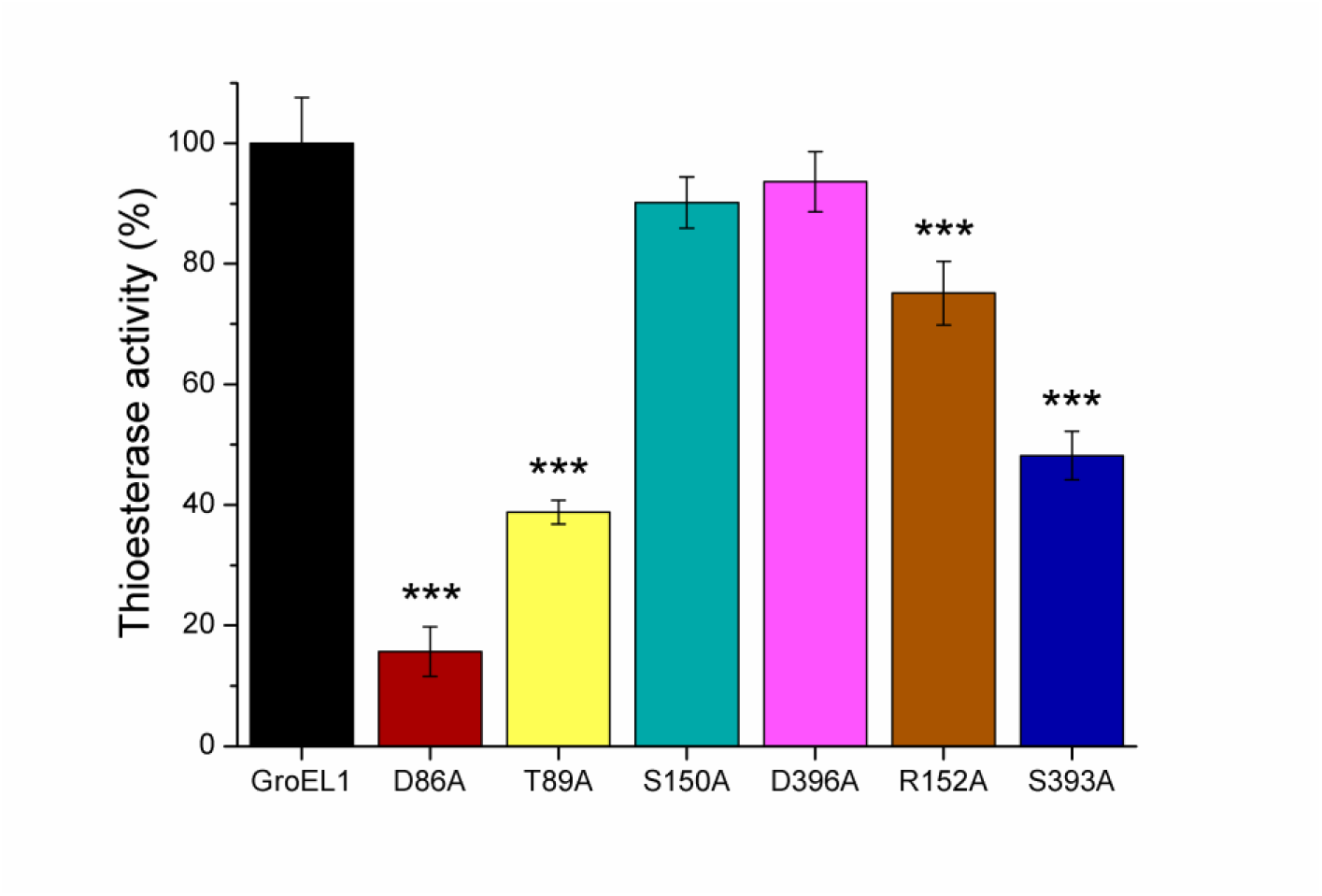
The thioesterase activity of GroEL1 mutant proteins in the presence of palmitoyl-CoA. The activity was monitored by measuring the absorbance at 412 nm for 1 h at 37 °C. The thioesterase activity of the D86A, T89A, S150A, D396A, R152A, S393A GroEL1 mutant proteins in comparison to the WT GroEL1 thioesterase activity. The mean from at least three independent experiments was calculated and plotted along with the standard deviation. The data were analyzed by two-sample unpaired *t* test: \*\*\**p*<0.001.

