## Supplementary Figures and Tables for "New Swiss-knife activities of GroEL/Hsp60 proteins"

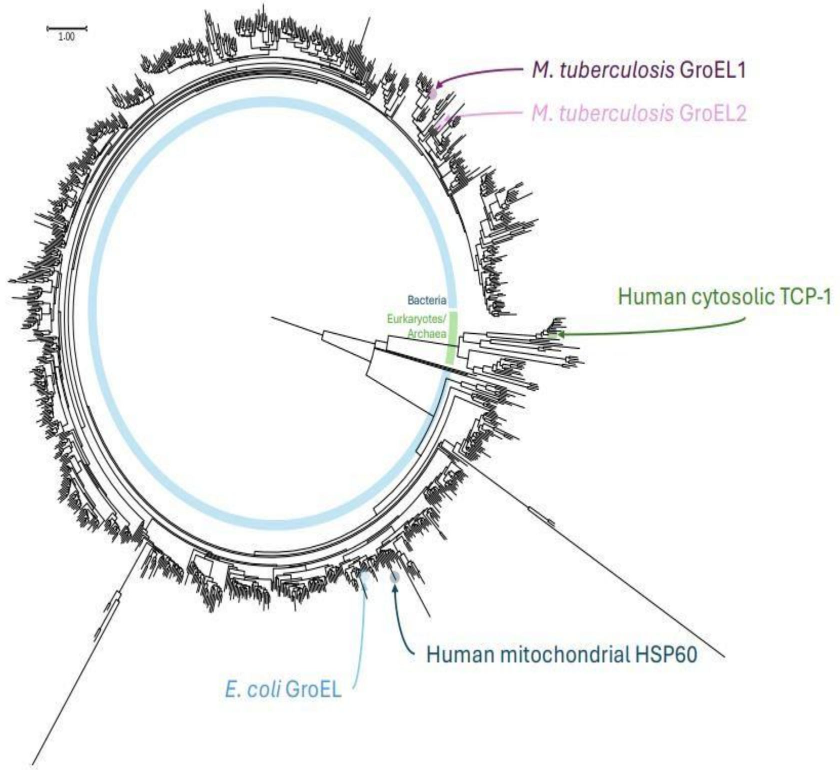


**Figure S1**: Pruned phylogenetic tree of GroEL proteins and their homologs.

**Table S1**. The primers for *pMT*GroEL and *pMT*Hsp60 plasmid constructions

| Oligonucleotides | Sequence (5' to 3') | Characteristics | Plasmid |
|---|---|---|---|
| **Hsp60-Fw** | 5'-GGAATCCATATGCTTCGGTTACCCACAGTC-3' | Nde1-Xho1 | pET-15b |
| **Hsp60-Rv** | 5'-GATATCCTCGAGTTAGAACATGCCACCTCCCAT-3' | Nde1-Xho1 | pET-15b |
| **GroEL-Fw** | 5'-GGAATCCATATGGCAGCTAAAGACGTA-3' | Nde1 | pET-15b |
| **GroEL-Rv** | 5'-GATATCCATATGTTACATCATGCCGCCCAT-3' | Nde1 | pET-15b |

**Table S2**. The primers for QuikChange Lightning Site-Directed Mutagenesis.

| mutation site | Primer Name | Primer Sequence (5' to 3’) |
|---|---|---|
| **Asp 86** | a257c_sense | 5'-atgtggccggtgccggcaccaccac-3' |
| | a257c_antisense | 5'-gtggtggtgccggcaccggccacat-3’ |
| **Thr 89** | a265g_sense | 5'-ggtgacggcaccgccaccgcaacca-3' |
| | a265g_antisense | 5'-tggttgcggtggcggtgccgtcacc-3’ |
| **Ser 150** | t448g_sense | 5'-tggcgacggtggcctcgcgcgac-3' |
| | t448g_antisense | 5'-gtcgcgcgaggccaccgtcgcca-3’ |
| **Asp 396** | a1187c_sense | 5'-aaagcgtcgaggctgcggtcgcggc-3' |
| | a1187c_antisense | 5'-gccgcgaccgcagcctcgacgcttt-3’ |
| **Arg 152** | c454g_g455c_sense | 5'-gacggtgtcctcggccgacgagcagatc-3' |
| | c454g_g455c_antisense | 5'-gatctgctcgtcggccgaggacaccgtc-3' |
| **Ser 393** | a1177g_g1178c_sense | 5'-cactcaaggagcgcaaggaagccgtcgaggatgc-3' |
| | a1177g_g1178c_ antisense | 5'-gcatcctcgacggcttccttgcgctccttgagtg-3' |

**Table S3**. The primers for the *pMT*PpsE-C plasmid construction

| Primer Name | Primer Sequence (5' to 3’) |
|---|---|
| **PP sense** | 5'-GGAATCCATATGCAGACCGAGGTTACGCTGCAA-3' |
| **COND antisense** | 5'-GGAATCCTCGAGATCTCACGTCGTGAACCAGCC-3' |
| **COND sense** | 5'-GGACTCCATATGAACGTCACGTACTTCCTCGAC-3' |
| **NRPS antisense** | 5'-GGAATCCTCGAGATCTCAGTCGTCGTGCTCTGT-3' |
| **END sense** | 5'-CACAACACTCTTAAGTCCGTTCCGCAC-3' |
| **END antisense** | 5'-GAATCCTCGAGTCAGCCGGCATCCAT-3' |

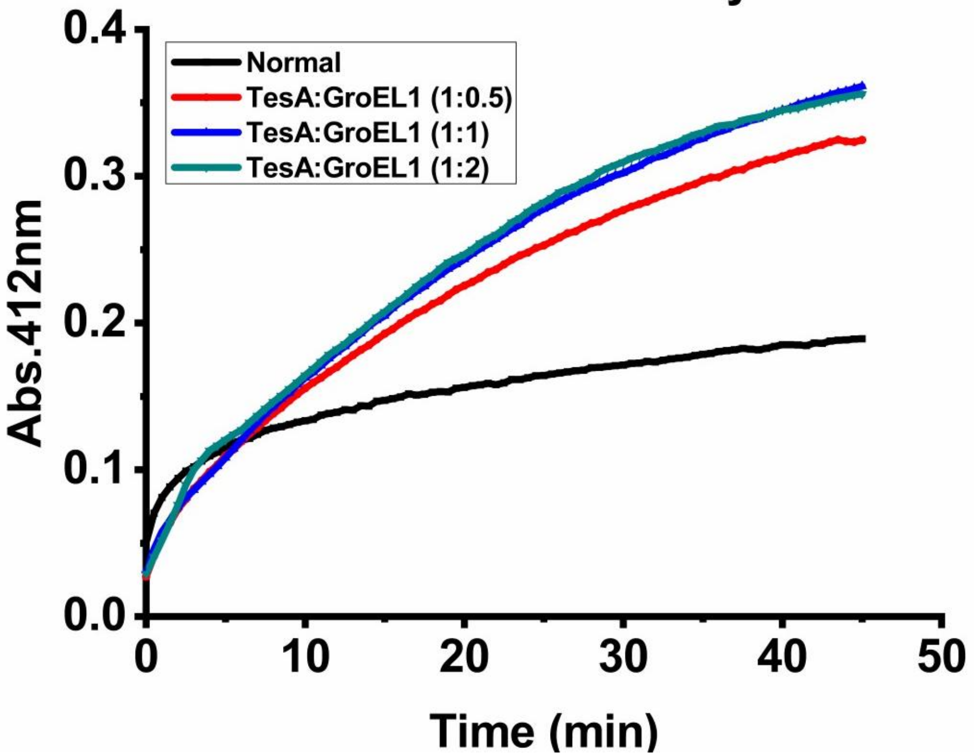


**Figure S2**. Effect of GroEL1 on TesA thioesterase activity. The reaction (200 μL) contained different protein concentration ratios, TesA : GroEL1 (1:0.5)/ (1:1)/ (1:2), 75 μM palmitoyl-CoA, 2.5 mM DTNB. Each plot is representative of three independent experiments.

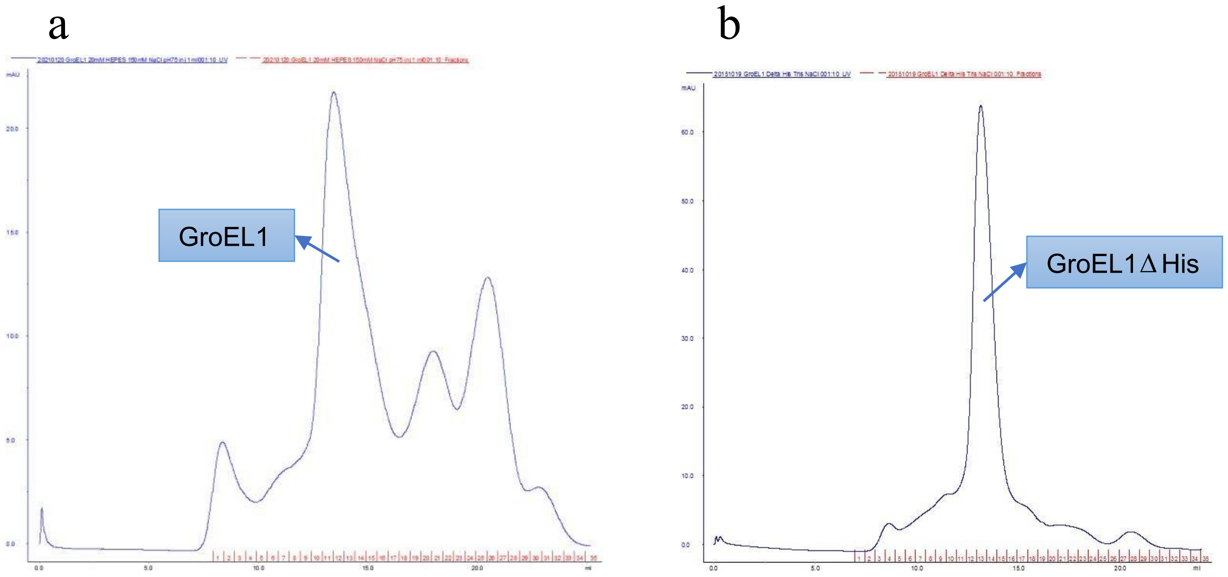


Figure S3A

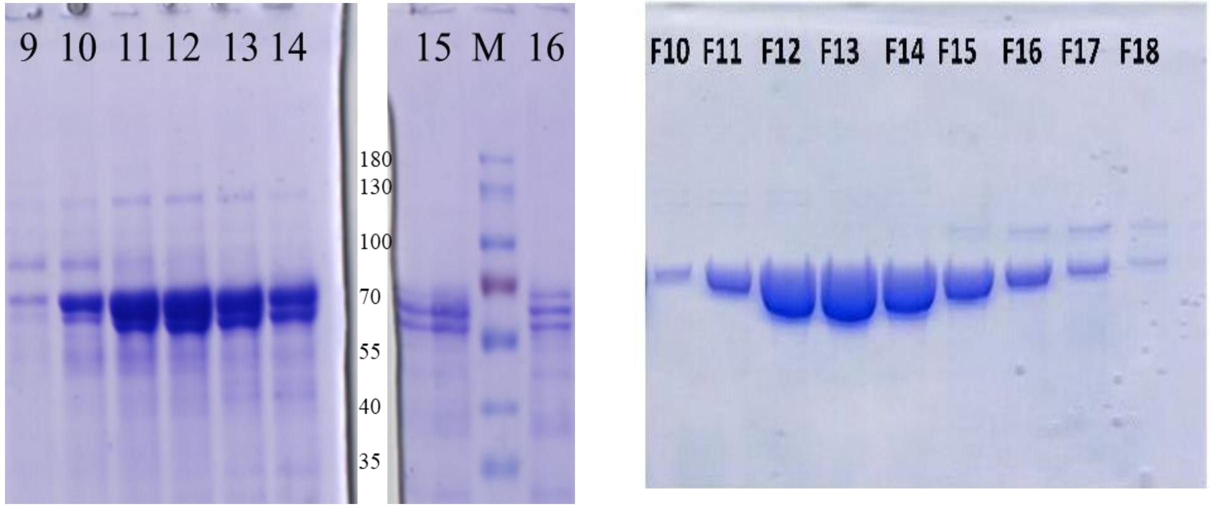


GroEL1 GroEL1 ΔHis

Figure S3B

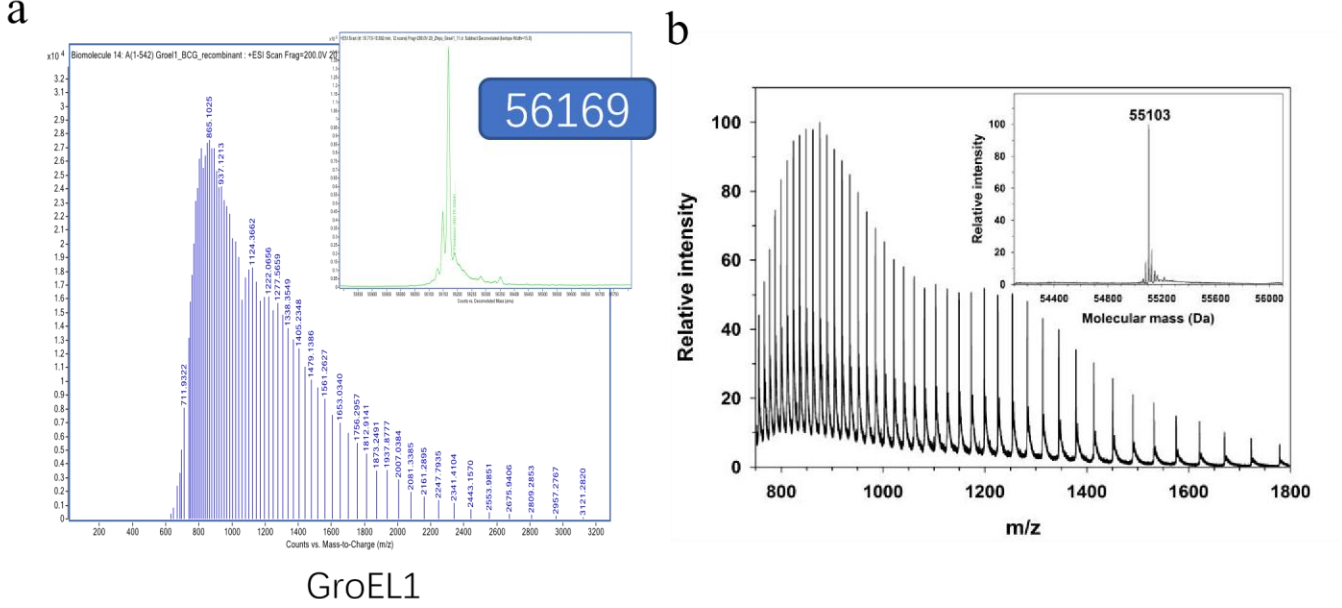


Figure S3C

**Figure S3**. Recombinant mycobacteria GroEL1 purity and integrity determination. (A) Size exclusion chromatography fractions: SEC elution profile of (a) recombinant GroEL1; (b) recombinant GroEL1 ΔHis. (B) Recombinant proteins purity determination by 12 % SDS-PAGE of fractions from the SEC: (a) recombinant GroEL1; (b) recombinant GroEL1 ΔHis. Lane M: PageRuler prestained protein ladder (Thermo Scientific). Fraction numbers of SEC eluate are shown on the top of the gels. Those figures are representative from at least three independent experiments. (C) Denaturing ESI mass spectrum of purified recombinant proteins: (a) recombinant GroEL1; (b) recombinant GroEL1 ΔHis. The experimental value is agreement with the theoretical molecular mass.

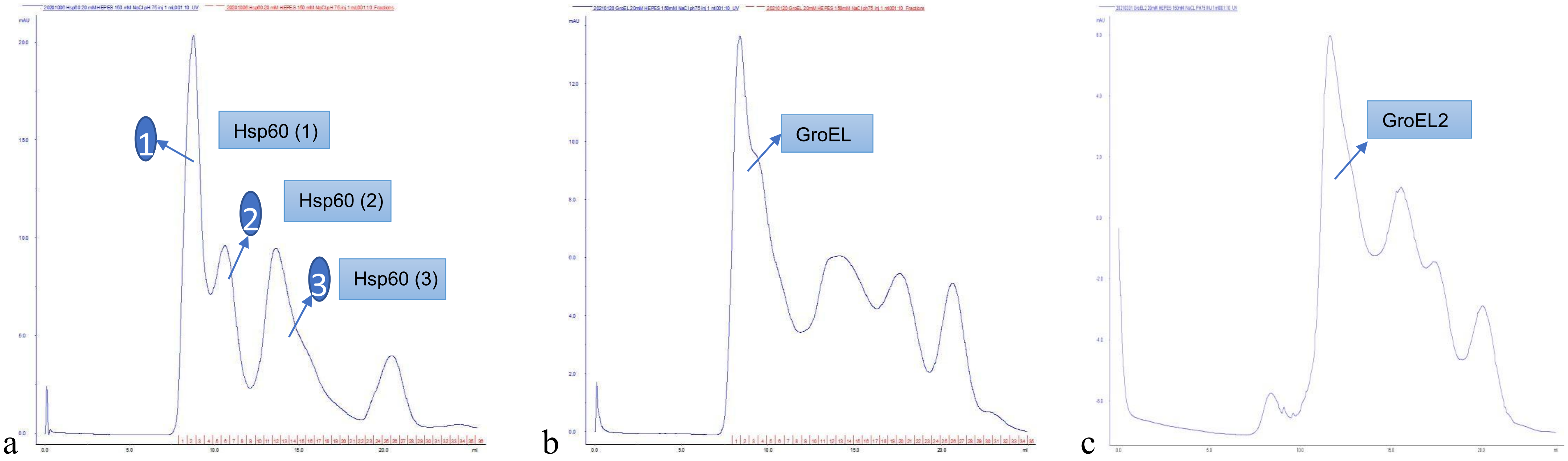


Figure S4A

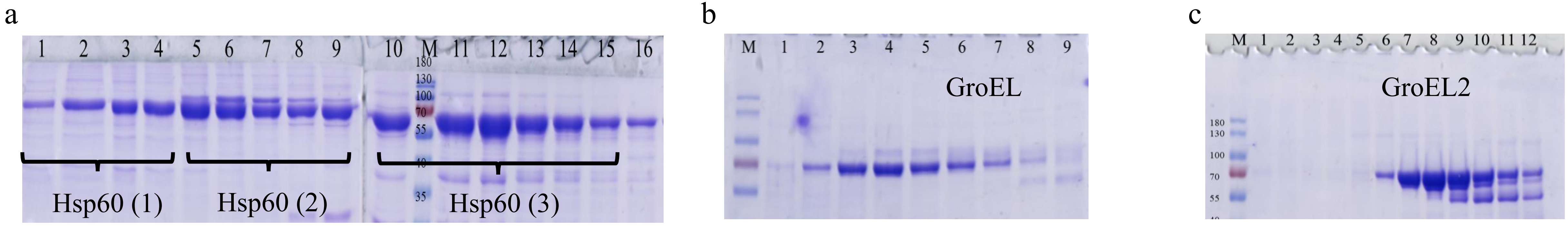


Figure S4B

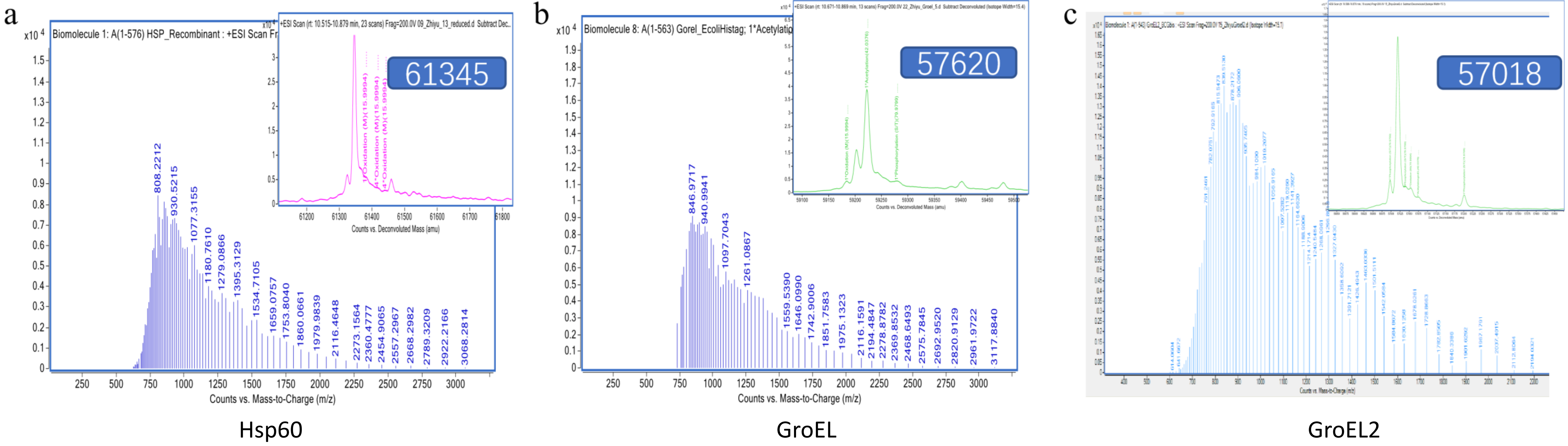


Figure S4C

**Figure S4**. Recombinant protein purity and integrity determination. (A) SEC elution profile of recombinant proteins: (a) human Hsp60; (b) *E.coli* GroEL; (c) *M. tuberculosis* GroEL2. (B) SEC fraction protein purity assessment by SDS-PAGE: (a) human Hsp60; (b) *E.coli* GroEL; (c) *M. tuberculosis* GroEL2. The figures are representative from at least three independent experiments. (C) Denaturing ESI mass spectrum of purified proteins: (a) Hsp60; (b) GroEL; (c) GroEL2. The experimental values are agreement with the theoretical molecular masses.

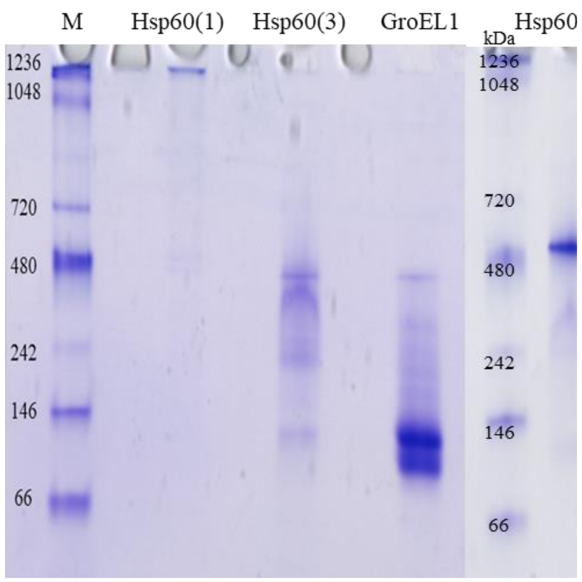


Figure S5A

| | Detection of various sub-units |
|---|---|
| GroEL | 14 |
| GroEL1 | 2, (7) |
| GroEL2 | 2-7 |
| Hsp60 (1) | 14 |
| Hsp60 (2) | 7 |
| Hsp60 (3) | 2-7 |

Figure S5B

**Figure S5**. (A) Oligomeric form assessment of the recombinant proteins or protein eluted fractions (Hsp60(1),(2),(3)) after SEC (Figure S1A and S3A) as determined by 4-15 % native-PAGE. (B) Main oligomeric forms of the recombinant proteins detected on the native gel. The number of subunits between brackets has been barely detected.

13

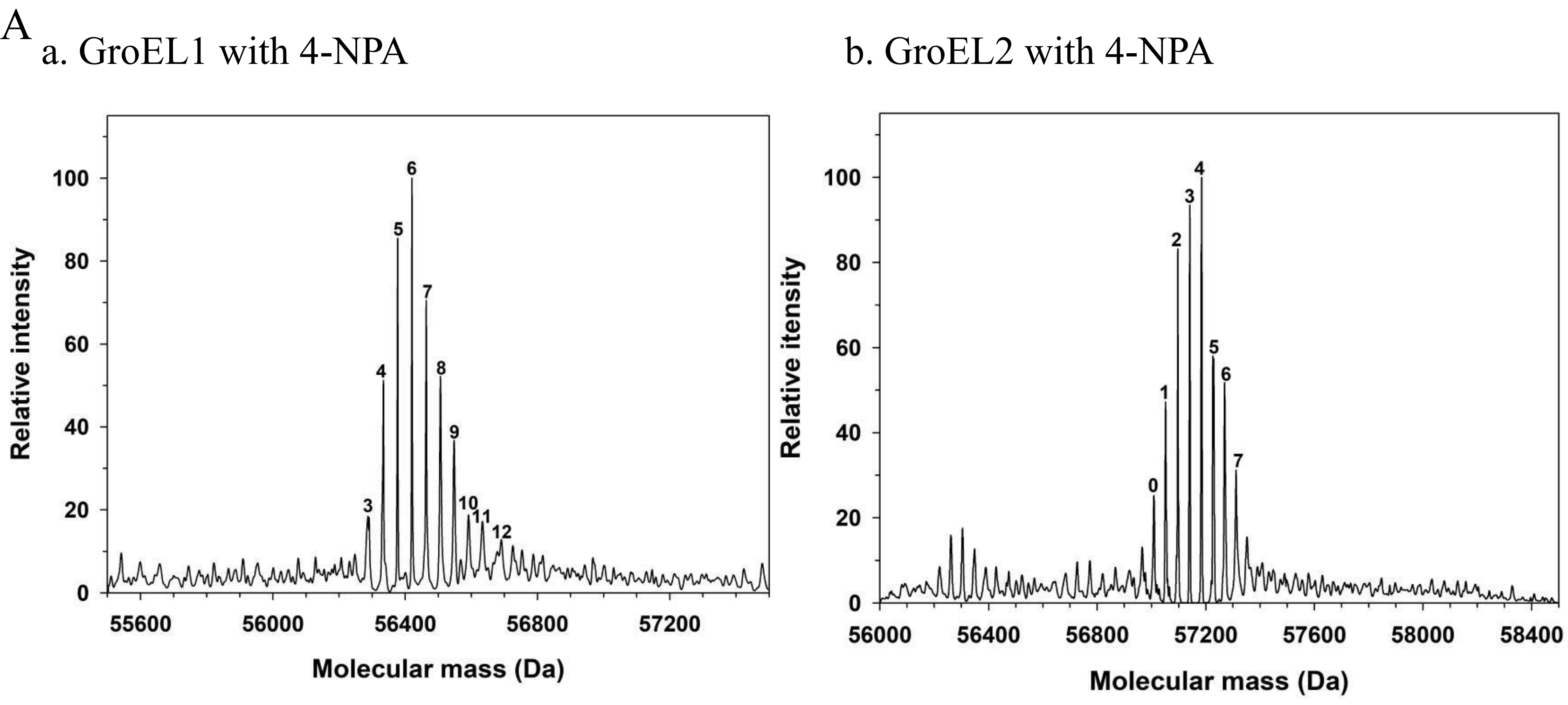


Figure S6A

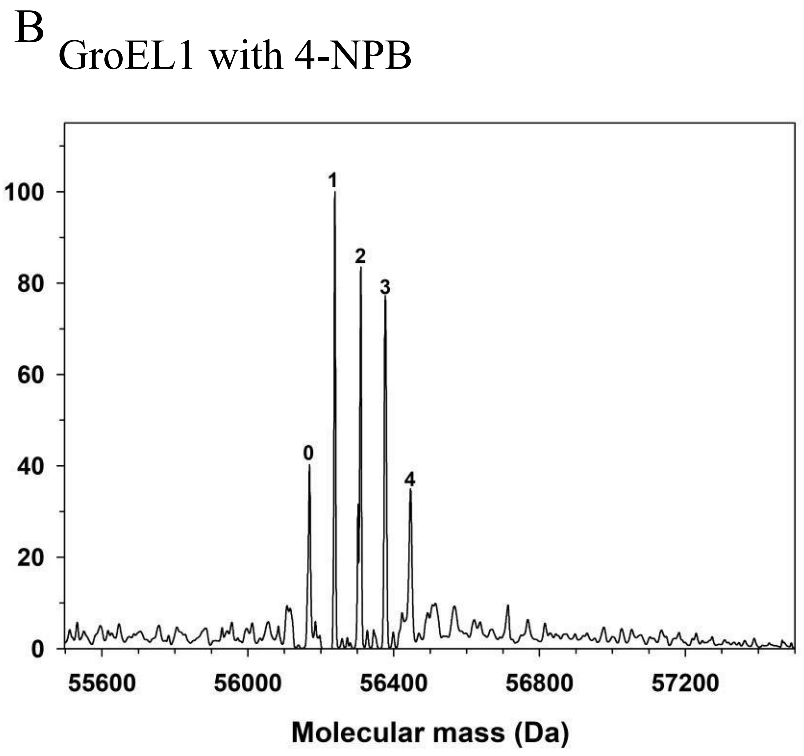


Figure S6B

C

```
GroEL1    MSKLIEYDETARRAMEVGMD KLADTVRVTLGPRGRHVVLA KAFGGPTVTNDGVTVAREIE LEDPFEDLGAQLVKSVATKT  80
GroEL2    MAKTIAYDEEARRGLERGLN ALADAVKVTLGPKGRNVVLE KKWGAPTITNDGVSIAKEIE LEDPYEKIGAELVKEVAKKT  80

GroEL1    NDVAGDGTTTATILAQALIK GGLRLVAAGVNPIALGVGIG KAADAVSEALLASATPVSGK TGIAQVATVSSRDEQIGDLV  160
GroEL2    DDVAGDGTTTATVLAQALVR EGLRNVAAGANPLGLKRGIE KAVEKVTETLLKGAKEVETK EQIAATAAISAGDQSIGDLI  160

GroEL1    GEAMSKVGHDGVVSVEESST LGTELEFTEGIGFDKGFLSA YFVTDFDNQQAVLEDALILL HQDKISSLPDLLPLLEKVAG  240
GroEL2    AEAMDKVGNEGVITVEESNT FGLQLELTEGMRFDKGYISG YFVTDPERQEAVLEDPYILL VSSKVSTVKDLLPLLEKVIG  240

GroEL1    TGKPLLIVAEDVEGEALATL VVNAIRKTLKAVAVKGPYFG DRRKAFLEDLAVVTGGQVVN PDAGMVLREVGLEVLGSARR  320
GroEL2    AGKPLLIIAEDVEGEALSTL VVNKIRGTFKSVAVKAPGFG DRRKAMLQDMAILTGGQVIS EEVGLTLENADLSLLGKARK 320

GroEL1    VVVSKDDTVIVDGGGTAEAV ANRAKHLRAEIDKSDSDWDR EKLGERLAKLAGGVAVIKVG AATETALKERKESVEDAVAA  400
GroEL2    VVVTKDETTIVEGAGDTDAI AGRVAQIRQEIENSDSDYDR EKLQERLAKLAGGVAVIKAG AATEVELKERKHRIEDAVRN  400

GroEL1    AKAAVEEGIVPGGGASLIHQ ARKALTELRASLTGDEVLGV DVFSEALAAPLFWIAANAGL DGSVVVNKVSELPAGHGLNV  480
GroEL2    AKAAVEEGIVAGGGVTLL-Q AAPTLDELK—LEGDEATGA NIVKVALEAPLKQIAFNSGL EPGVVAEKVRNLPAGHGLNA  477

GroEL1    NTLSYGDLAADGVIDPVKVT RSAVLNASSVARMVLTTETV VVDKPAKAEDHDHHHGHAH---- 539
GroEL2    QTGVYEDLLAAGVADPVKVT RSALQNAASIAGLFLTTEAV VADKPEKEKASVPGGGDMGGMDF  540
```

Figure S6C

**Figure S6**. (A) Analysis by mass spectrometry of recombinant mycobacteria protein acetylation in the presence of 4-NPA. (a) GroEL1; (b) GroEL2. The number above each peak represents the amount of acetylation. (B) Analysis of recombinant GroEL1 butyrylation in the presence of 4-NPB by mass spectrometry. The number above each peak represents the amount of acetylation. (C) Protein sequence alignment of recombinant GroEL1 and GroEL2 with acetylated lysine residues indicated in red in GroEL1 and in blue in GroEL2. The other acetylated residues are highlighted in yellow. The acetylated lysine residues identified by Xie *et al.* in 2015 and Birhanu *et al.* in 2017, are underlined in red and highlighted in green, respectively.

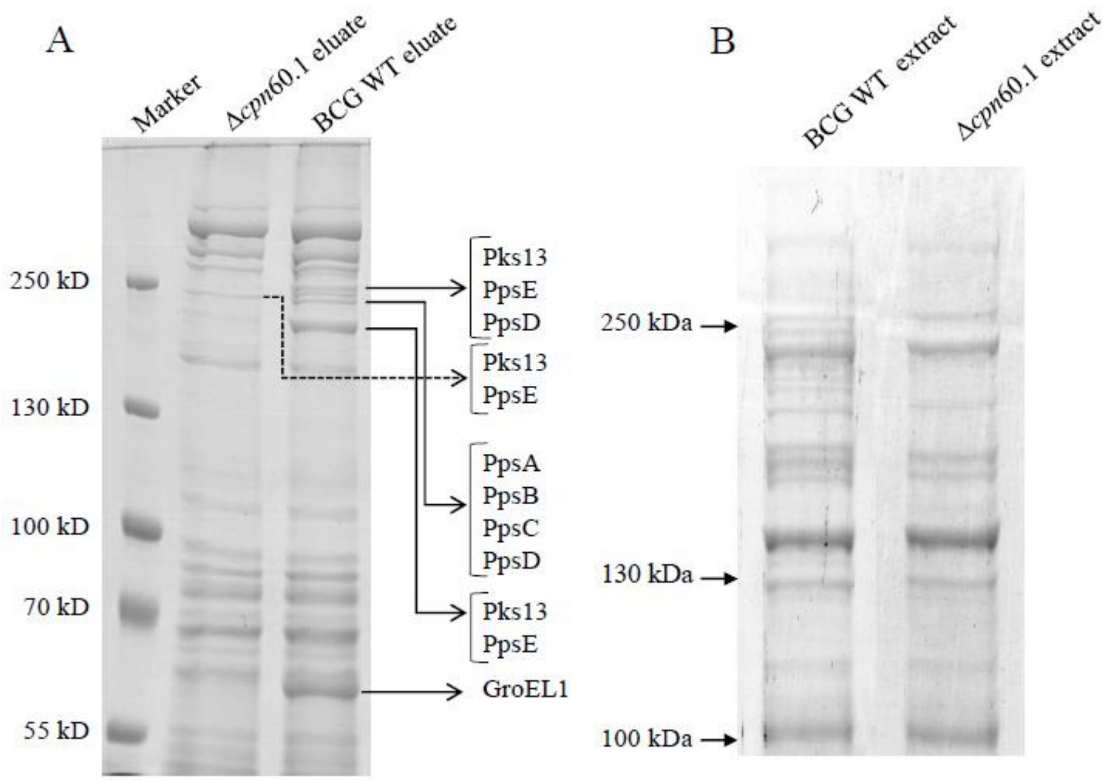


**Figure S7.** Co-elution of GroEL1 and PKS proteins from the WT and Δ*cpn*60.1 *M. bovis* BCG. (A) Protein eluates after Ni-column purification were analyzed by 12 % SDS-PAGE, and further characterized by MS/MS after gel elution and trypsin digestion. (B) Protein extracted from the WT and Δ*cpn*60.1 *M. bovis* BCG according to 12 % SDS-PAGE analysis.

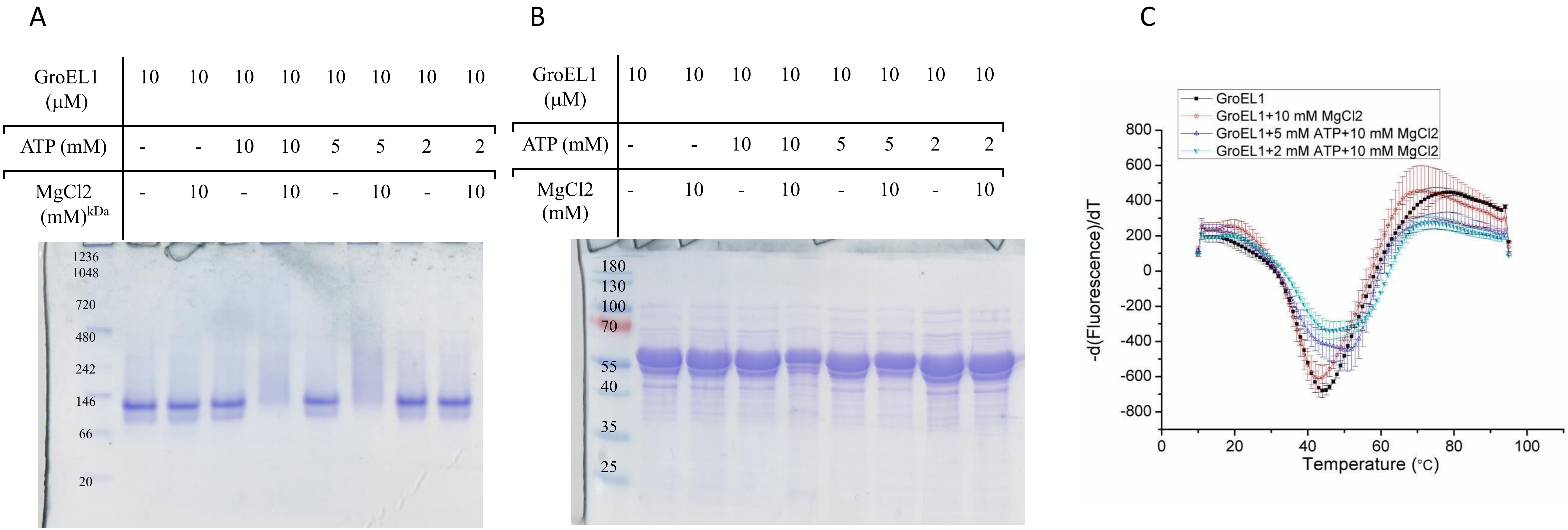


**Figure S8**. (A) The impact of ATP and Mg on *M. tuberculosis* GroEL1 oligomerization. (a) GroEL1 dimers are not detected on native polyacrylamide gel in the presence of ATP and Mg. (b) GroEL1 could be detected in denaturing condition in the presence of ATP and Mg. (B) Thermal shift assay analysis of the effect of ATP and Mg on GroEL1 stability. The reaction (25 μL) contained 5 μM GroEL1, 2.5 × SYPRO Orange, with 5 mM /2 mM ATP and 10 mM $MgCL_2$ in 20 mM HEPES, 150 mM NaCL, pH 7.5. Those figures are representative from at least three independent experiments.

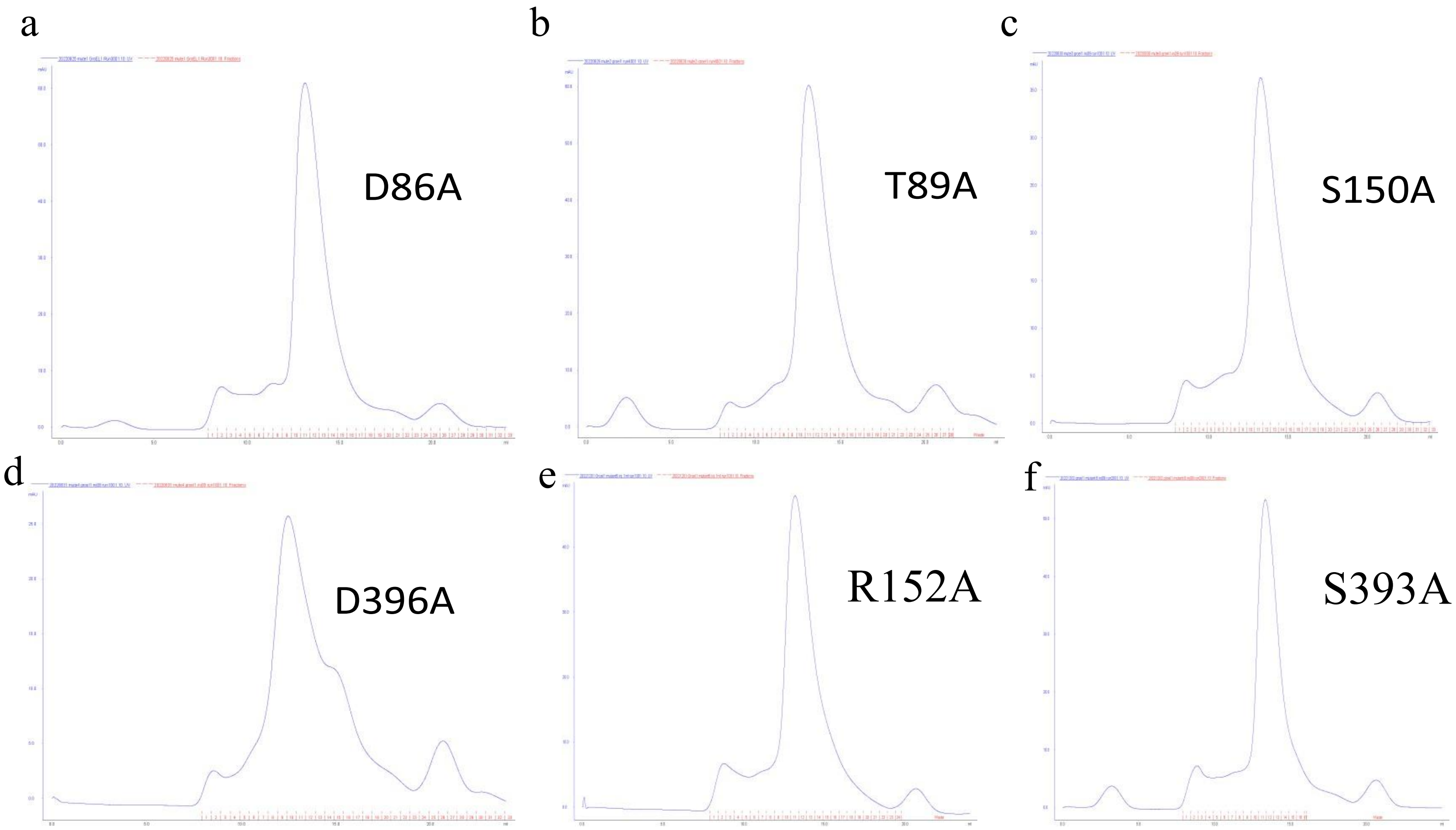


Figure S9A

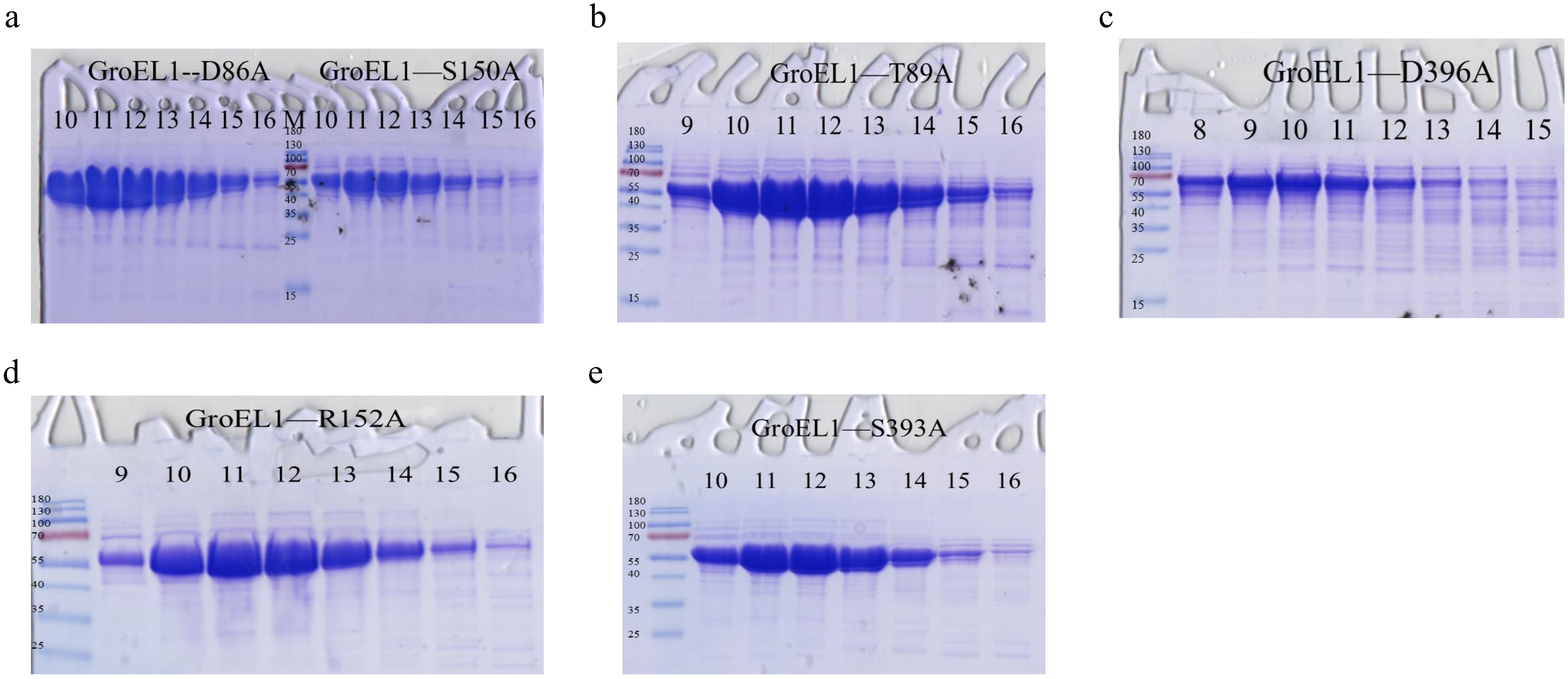
a
GroEL1--D86A
GroEL1—S150A
10 11 12 13 14 15 16 M 10 11 12 13 14 15 16
b
GroEL1—T89A
9 10 11 12 13 14 15 16
c
GroEL1—D396A
8 9 10 11 12 13 14 15
d
GroEL1—R152A
9 10 11 12 13 14 15 16
e
GroEL1—S393A
10 11 12 13 14 15 16

Figure S9B

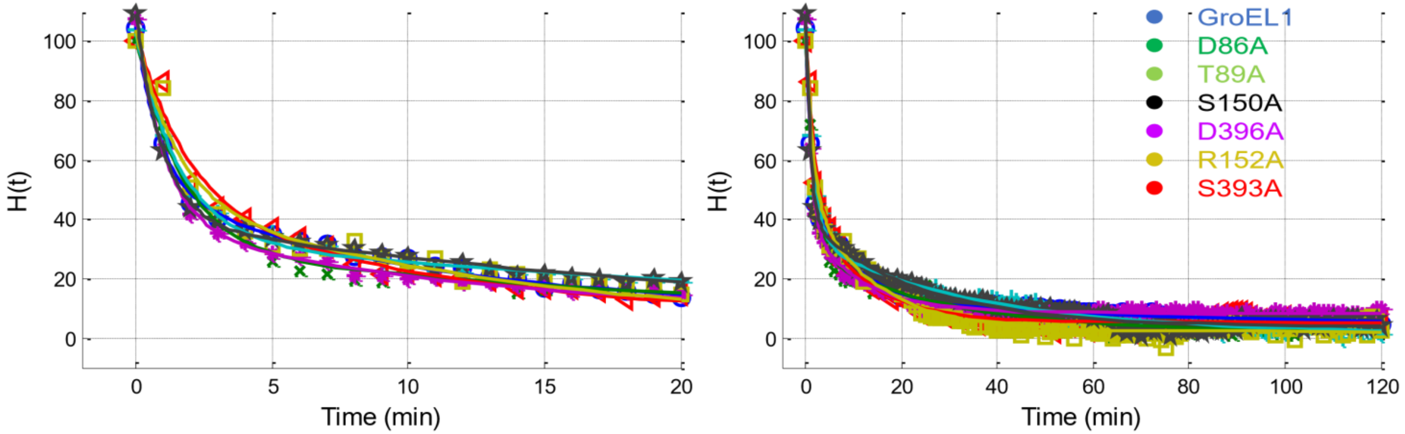


Figure S9C

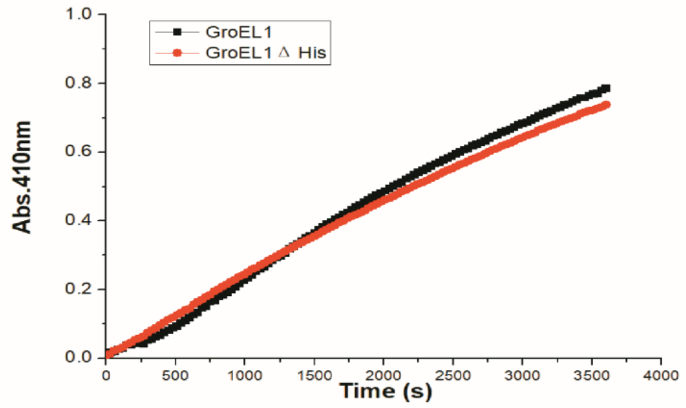


Figure S9D

**Figure S9**. Six different recombinant mutagenized GroEL1 purity determination. (A) Size exclusion chromatography fractions: SEC elution profile of recombinant mutant GroEL1 (a) D86A; (b) T89A; (c) S150A; (d) D396A; (e) R152A; (f) S393A. (B) Recombinant proteins purity determination by 12 % SDS-PAGE of fractions from the SEC: recombinant mutant GroEL1 (a) D86A and S150A; (b) T89A; (c) D396A; (d) R152A; (e) S393A. Lane M: PageRuler prestained protein ladder (Thermo Scientific). Fraction numbers of SEC eluate are shown on the top of the gels. (C) FTIR analysis of the tertiary structure of purified mutant proteins. Those figures are representative from at least three independent experiments. (D) Hydrolytic activities of GroEL1 and GroEL1 ΔHis.

**Table S4**. Residues mutated in GroEL1 mapped via the global MSA onto corresponding residues in other proteins of interest.

| Logo | Pos. MSA | GroEL | | GroEL1 | | GroEL2 | | HSP 60 | | TCP-1 | |
|---|---|---|---|---|---|---|---|---|---|---|---|
| 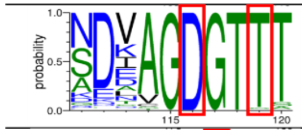 | 116 | 87 | **D** | 86 | **D** | 86 | **D** | 111 | **D** | 88 | **D** |
| | 119 | 90 | **T** | 89 | **T** | 89 | **T** | 114 | **T** | 91 | **T** |
| 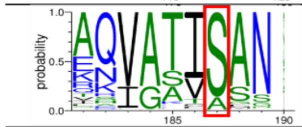 | 187 | 151 | **S** | 150 | **S** | 150 | **S** | 175 | **S** | 155 | **S** |
| 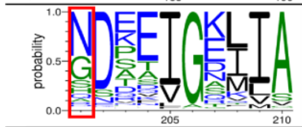 | 201 | 154 | S | 152 | R | 152 | G | 178 | G | 164 | **N** |
| 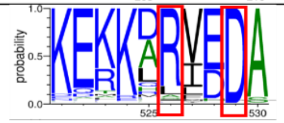 | 526 | 395 | R | 393 | S | 393 | R | 420 | R | 391 | S |
| | 529 | 398 | **D** | 396 | **D** | 396 | **D** | 423 | **D** | 394 | **D** |
